# RAxML-NG 2: Automatic Model Selection, Novel Tree Search Heuristics, and Fast Branch Support Metrics

**DOI:** 10.64898/2026.09.09.750097

**Authors:** Oleksiy M. Kozlov, Anastasis Togkousidis, Christoph Stelz, Dimitri Höhler, Julius Wiegert, Alexandros Stamatakis

**Affiliations:** Computational Molecular Evolution Group, Heidelberg Institute for Theoretical Studies, Heidelberg, Germany; Biodiversity Computing Group, Institute of Computer Science, Foundation for Research and Technology - Hellas, Heraklion, Crete, Greece; Institute for Theoretical Informatics, Karlsruhe Institute of Technology, Karlsruhe, Germany; Hector Institute for Artificial Intelligence in Psychiatry, Central Institute of Mental Health, Medical Faculty Mannheim, Heidelberg University, Mannheim, Germany; Department of Psychiatry and Psychotherapy, Central Institute of Mental Health, Medical Faculty Mannheim, Heidelberg University, Mannheim, Germany

**Keywords:** Phylogenetic Inference, Model selection, Branch support, Maximum Likelihood

## Abstract

RAxML-NG is a widely used tool for maximum likelihood based phylogenetic inference. In the seven years since the last RAxML-NG publication, we have continuously improved and extended the code. Here, we describe the next major release, RAxML-NG 2.0. It introduces a number of new features: integrated model testing, multiple fast branch support metrics, automatic parallelization tuning, phylogenetic difficulty prediction, genotype evolution models, to name but the most important ones. Furthermore, we introduce two novel search heuristics at production code level: the adaptive difficulty-aware heuristic (default) and the fast mode with early-stopping that prevents over-optimization. We perform extensive benchmarking of RAxML-NG 2.0 with respect to its accuracy and speed, and compare it to other popular maximum likelihood based phylogenetic inference tools (IQTree, VeryFastTree) as well as to preceding RAxML-NG versions. In particular, the new fast search heuristic in conjunction with machine learning based branch support prediction induces a 65*×* inference time reduction compared to RAxML-NG 1.2, with minor to no accuracy loss.

The code is available under GNU GPL at https://codeberg.org/amkozlov/raxml-ng.

## Introduction

RAxML-NG (1) is one of the most widely used tools for maximum likelihood (ML) based phylogenetic inference, along with IQTree (2), RAxML (3), PhyML (4), and (Very)FastTree (5, 6). It has been used in numerous high-profile studies to analyze viral (7), microbial (8), avian (9), plant (10), and human (11) datasets, for instance.

Over the past seven years, we have continuously improved RAxML-NG by incorporating novel algorithms and models as well as user feedback. In this new release, we implement automatic model selection (MOOSE), which will help to substantially simplify and expedite typical phylogenetic workflows. Furthermore, we address the arguably most important inefficiency in RAxML-NG 1.x, that is, the slow branch support estimation via the standard Felsenstein bootstrap procedure (12). RAxML-NG 2.0 now implements multiple of well-established fast support metrics such as the rapid boot-strap and SH-aLRT (Shimodaira-Hasegawa-like approximate Likelihood Ratio Test), as well as our recently introduced machine learning based support value predictor called EBG (Educated Bootstrap Guesser, Table 1).

**Table 1.** Branch support metrics available in RAxML-NG 2.

| Abbr. | Name | Reference |
|---|---|---|
| FBP | Felsenstein Bootstrap | (12) |
| TBE | Transfer Bootstrap Expectation | (17) |
| RBS | Rapid Bootstrap (RAxML8-style) | (18) |
| EBG | Educated Bootstrap Guesser<sup>*a*</sup> | (19) |
| TCF | Tree Concordance Factor<sup>*b*</sup> | (20) |
| SH | SH-like aLRT | (4) |
| IC | Internode Certainty | (21) |
| PS | Parsimony Support | (19) |
| PBS | Parsimony Bootstrap Support | (12) |

<sup>*a*</sup> EBG-Light with a reduced set of features (see Supplementary methods: EBG-Light)
<sup>*b*</sup> Identical to gCF (22) for taxon-complete trees (see Supplementary methods: Tree Concordance Factor).

## Materials and Methods

### New Features and Improvements

#### Adaptive search heuristics

RAxML-NG 1.1 relies upon a thorough tree search heuristic that generates 20 starting trees by default (10 random + 10 parsimony-based) to conduct 20 independent ML tree searches. This method was optimized to find the tree with the best possible ML score on most datasets, yet its computational cost can be excessively high. Based upon a systematic exploration of alternative search algorithm configurations, we found that obtained ML score improvements are often statistically insignificant. Hence, investing additional optimization effort and CPU cycles does not necessarily yield a qualitatively as well as quantitatively improved tree topology. Based on these findings, we relaxed several convergence thresholds in RAxML-NG 1.2 to accelerate tree searches by up to 1.9× (13).

In RAxML-NG 2.0, we deploy an adaptive tree search strategy (14) that fine-tunes the number of starting trees as well as the search heuristic parameters (e.g., the Subtree Pruning and Re-grafting (SPR) radius), based on the Pythia difficulty score (15) for the specific dataset at hand. More specifically, we observed that on *easy* MSAs there is typically a single likelihood peak, which can be found by a superficial search from a single starting tree. Conversely, on *difficult* MSAs, multiple tree searches converge to a topologically distinct tree; however, the differences between their likelihood scores are usually statistically insignificant. Finally, on MSAs with *medium* difficulty, there are often several clear likelihood peaks, and hence multiple thorough searches from distinct starting trees help to escape local optima and yield a significantly better tree. Based on these observations, we devised an adaptive search heuristic which maximizes the search effort for the datasets which are likely to benefit from it, and progressively reduces search effort for less promising datasets.

This adaptive heuristic is now the *default* tree search method in RAxML-NG 2.0 (Figure 1a).

**Fig. 1.**
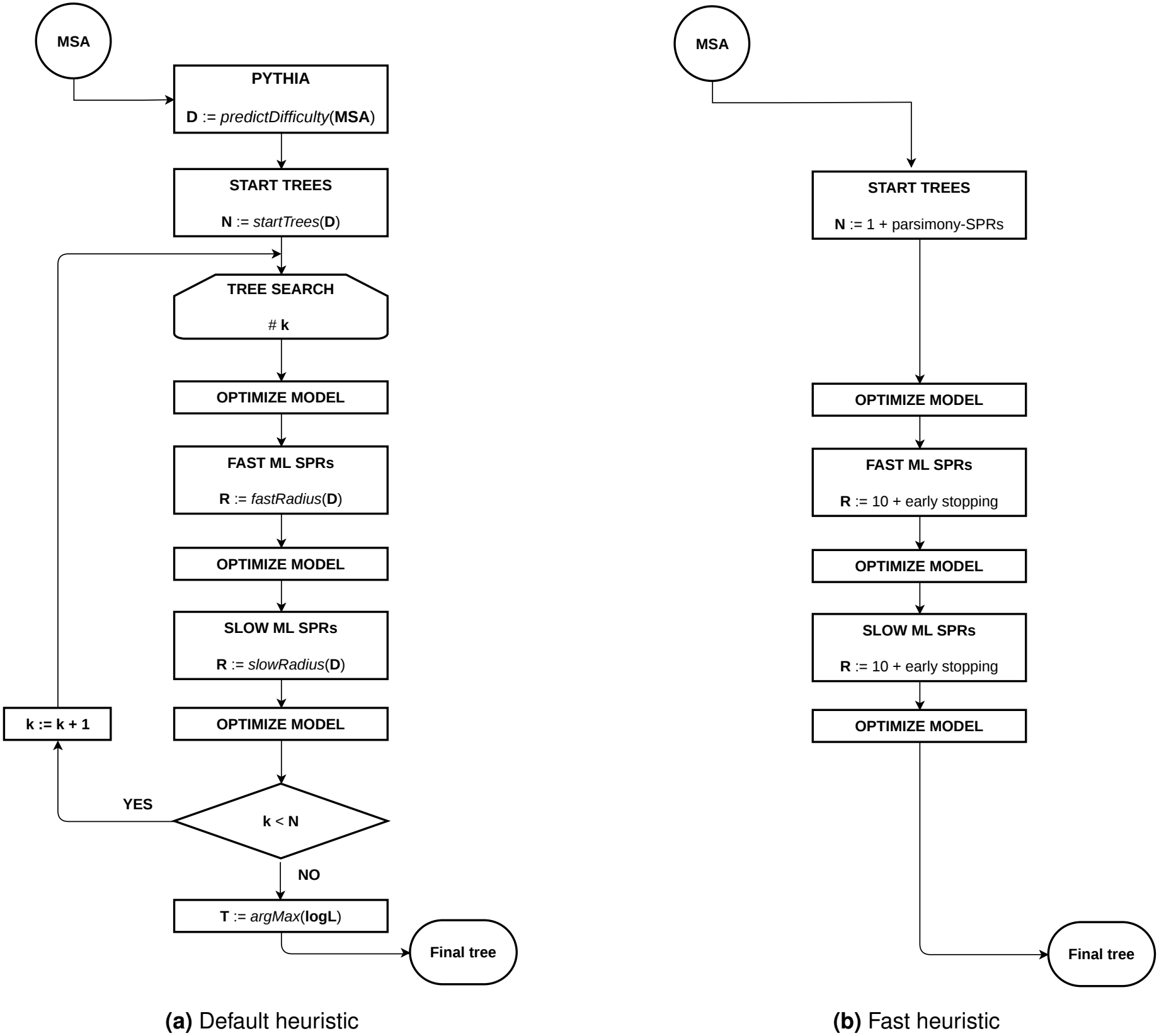
Simplified flowcharts of the default *(a)* and fast *(b)* tree search heuristics in RAxML-NG 2. The default heuristic conducts multiple searches from both parsimony-based and random starting trees. The number thereof *N* and the radius of the SPR moves (*R*) are determined by the predicted MSA difficulty (*D*). A tree with the highest log-likelihood score across all *N* searches is selected as the final tree *T*. The fast heuristic performs a single tree search from a parsimony-based starting tree. Both fast and slow SPR move rounds have a fixed re-grafting radius of 10. An early stopping heuristic based on the KH-test is applied to reduce the number of SPR rounds.

In this paper, we also introduce a novel *fast* search heuristic (Figure 1b) that performs a limited number of SPRs with a fixed re-grafting radius of 10 that starts from a single parsimony tree. In the fast search mode, we enable the Kishino–Hasegawa (KH) test to detect and circumvent overoptimization (16).

We provide a detailed description of both *default* and *fast* search heuristics in the Supplementary methods: Tree search heuristics.

#### Fast branch support metrics

To infer branch support values, RAxML-NG 1.x uses the standard non-parametric bootstrap method proposed by Felsenstein (FBP). The FBP, although widely accepted, requires a full ML tree search on every bootstrap replicate, which yields it computationally intractable for large datasets. Hence, in RAxML-NG 2.0, we implement a plethora of alternative, faster, and novel branch support metrics (Table 1). First, we re-implement the rapid bootstrap algorithm (RBS) from RAxML 8 (18). RBS is substantially faster than FBP due to a more superficial ML tree search heuristic. Nonetheless, it approximates FBP supports reasonably well, as has been recently confirmed (19). An alternative approach, the Educated Bootstrap Guesser (19), uses a LightBGM model (23) to predict support values from a set of (comparably) inexpensive-to-compute dataset features. By avoiding expensive ML searches, it exhibits superior speed as well as accuracy compared to RBS, UF-BOOT2 (24), and other state-of-the-art support metrics (19). In RAxML-NG 2.0, we implement a simplified EBG version that relies upon a reduced feature set (see Supplementary methods: EBG-Light for details).

#### Automatic multi-grain parallelization

RAxML-NG supports both, *fine-grained* parallelization across alignment patterns, as well as *coarse-grained* parallelization across multiple starting trees or bootstrap replicates. In fine-grained parallelization mode, the number of CPU cores (threads) that can be efficiently utilized is limited by the number of alignment patterns. Conversely, the scalability of the coarse-grain parallelization mode is limited by the number of starting trees or bootstrap replicates. RAxML-NG uses a simple, fast-to-compute heuristic to estimate the optimal number of alignment patterns per thread for the dataset at hand and subsequently automatically configure the multi-grain parallelization scheme accordingly. On very small datasets (w.r.t. number of alignment patterns), we reduce the number of threads to improve efficiency and save energy. Users can override this automatic parallelization scheme by explicitly specifying the number of threads and parallel tree searches.

#### Integrated model selection

Automatic model testing is being routinely conducted prior to phylogenetic tree inference to determine the best-fit evolutionary model for the MSA at hand. Multiple stand-alone tools and pipelines exist for this task, such as, ModelTest-NG (25) or ParGenes (26). We now integrate the model selection procedure directly into RAxML-NG 2 to improve user experience and simplify typical workflows. Like IQ-TREE’s ModelFinder (27), it supports FreeRate models and uses heuristics to reduce the set of candidate models. MOOSE benefits from the mixed fine- and coarse-grained parallelization described above: It is able to evaluate multiple candidate models concurrently, thereby reducing the need of synchronization and increasing the utilization of modern multi-core processors. Since the aforementioned candidate selection heuristics introduce dependencies between models, we use model prioritization and speculative model evaluation to improve parallel efficiency. The details of this model selection algorithm are described in the Supplementary methods.

#### Genotype models and VCF support

For diploid organisms, their genotype can be encoded as nucleotide pair, that is, an allele for each chromosome. If the chromosome assignment is known (“phased”), the pair is ordered and hence there exist 16 possible genotype states. Otherwise, we consider an “unphased” genotype with 10 distinct unordered nucleotide pairs.

In RAxML-NG 2.0, we integrate substitution models for phased (GT16) and unphased (GT10) diploid genotype data originally introduced in CellPhy (28), a dedicated tool for single-cell phylogeny reconstruction. These models make two simplifying assumptions to improve computational tractability: (1) only one of the two alleles in a genotype can change in an infinitesimal amount of time, hence substitution rates that correspond to “double-mutations” are fixed to a very small value (e.g., *r*(*A*|*A* ↔ *T* |*T*) = 10^−6^), and (2) nucleotide substitution rates are independent of their allelic position (e.g., *r*(*A*|*C* ↔ *A*|*T*) = *r*(*C*|*G* ↔ *T* |*G*) = *r*(*C* ↔ *G*)). Genotype input data can be provided either as a FASTA or PHYLIP file with a dedicated character state encoding (see Table S2), or as a standard VCF file. In the latter case, RAxML-NG offers two modes. In *deterministic VCF* mode, it reads the GT field (that stores the genotype calls), and builds a classical, discrete alignment matrix with fixed genotype states. In *probabilistic VCF* mode, it reads the PL field (which stores the Phred-scaled genotype likelihoods) and uses the likelihood of each genotype to initialize the corresponding per-state probabilities at the tips of the tree thereby using a probabilistic alignment without discrete states.

For a detailed description of these genotype models, please refer to (28).

### Usage and availability

The source code of RAxMLNG is publicly available at https://codeberg.org/amkozlov/raxml-ng; we provide pre-compiled binaries for Linux and macOS with every release. Furthermore, RAxML-NG can be installed via popular package managers such as *bioconda, homebrew, EasyBuild*, and *SPACK*. Finally, RAxML-NG can be used without a local installation through the following web services: https://phylogeny.fr (basic) and https://usegalaxy.eu (advanced).

We validated the code quality of RAxML-NG 2.0.3 with SoftWipe (29) and obtained a score of 8.1 / 10, which is similar to the previous stable version (1.2.2).

### Experimental setup

#### Hardware

For tree search, branch support, and model selection benchmarks, we used a server with a dual-socket AMD EPYC 9684X CPU (192 physical cores). In all experiments, 64 CPU cores (1/3) remained idle to minimize the effects of thermal throttling and job interference.

#### Datasets

For the runtime and accuracy evaluation of RAxML-NG, we used empirical as well as simulated DNA and amino acid (AA) datasets.

First, we used the 222 empirical DNA, and 78 empirical AA datasets that were analyzed in Togkousidis et al. (16). The authors ranked TreeBASE (30) datasets based on their size (as quantified by number of taxa × number of sites) and selected the 300 largest datasets. We subsequently removed one DNA dataset since it could not be analyzed with IQ-TREE3, and included an additional 9 empirical AA alignments of large protein families that comprise a high number of taxa (698 − 4390) and exhibit high phylogenetic difficulty (0.50 − 0.77). Overall, we conducted the accuracy evaluation on 221 empirical DNA, and 87 empirical AA datasets.

Second, we simulated 150 DNA and 150 AA datasets based on trees and parameter distributions from our RAxML-Grove (31) database. RAxMLGrove contains anonymized trees inferred by RAxML and RAxML-NG users including the respective model parameter estimates. Since the Tree-BASE datasets are comparatively small (median: 226 taxa, 3401 site patterns in our benchmark subset), we simulated larger datasets with regard to the numbers of sites and taxa. The size dimensions were chosen such that the resulting MSAs were larger than the TreeBASE average, while keeping overall benchmark runtimes manageable (i.e., a maximum runtime for the slowest tools of around two weeks on our benchmarking server). Specifically, we used the following strategy to simulate MSAs:

1. Define target dataset dimensions as #taxa (*t*_*n*_) × #sites (*s*_*n*_): (DNA) *t*_*n*_ ∈ [900, 1600, 3600] × *s*_*n*_ ∈ [4000, 8000, 16000, 32000, 64000], (AA) *t*_*n*_ ∈ [450, 900, 1800] × *s*_*n*_ ∈ [500, 1000, 2000, 4000, 8000].
2. For each target dataset size, generate 10 replicate MSAs as follows:
  a. Select a dataset reference (i.e., inferred tree and estimated model parameters) from RAxMLGrove where *(1)* the number of taxa is up to twice our target value *t*_*n*_, and *(2)* inference model is GTR+Γ (DNA) or LG+Γ/LG+CAT (AA). We draw reference datasets randomly with replacement. Subsequently, we randomly prune taxa until the target tree size *t*_*n*_ is reached.
  b. Simulate MSAs with RAxMLGroveScripts (31) (RGS) (a custom AliSim (32) wrapper), using the empirical trees and model parameters from the respective RAxMLGrove reference datasets, and the desired number of sites *s*_*n*_. We did not use any insertion or deletion models for our simulation. However, for partitioned datasets RGS uses the available presence-absence-matrices (which encode 0 if a sequence is not present in a given partition, and 1 otherwise) to imitate the empirical missing-data patterns on partition level.

We removed all duplicate sequences from the empirical and simulated datasets prior to ML inference. Characteristics of all empirical and simulated datasets used for evaluation are summarized in Table 2.

**Table 2.** Datasets used for benchmarking.

| Source<sup>a</sup> | Datatype<sup>a</sup> | # taxa<sup>b</sup> | # sites | Difficulty |
|---|---|---|---|---|
| Empirical (308) | DNA (221) | 15 to 1,200 | 236 to 607,736 | 0.01 to 0.80 |
|  | AA (87) | 18 to 4,390 | 98 to 759,490 | 0.01 to 0.77 |
| Simulated (300) | DNA (150) | 249 to 3600 | 4000, 8000, 16000, 32000, 64000 | 0.02 to 0.82 |
|  | AA (150) | 217 to 1800 | 500, 1000, 2000, 4000, 8000 | 0.12 to 0.46 |

<sup>a</sup> Number of datasets is specified in parentheses.
<sup>b</sup> After the removal of duplicate sequences.

#### Software and metrics

To evaluate tree inference accuracy and speed, we used the PhyloSmew pipeline (33). In short, the pipeline works as follows. In a configuration file, we define the tools to be benchmarked (by specifying their respective command lines) and the respective benchmark datasets (see Table 2). Then, all tools are executed on all datasets. Once all inferences have been completed, the pipeline computes corresponding per-tool performance metrics such as the runtime, the Robinson-Foulds (RF) distance (34) to the reference tree, and the statistical plausibility using the Approximately Unbiased (AU) test (35) as implemented in CONSEL (36).

We use *percentage of plausible trees* as our primary accuracy metric. An inferred tree is considered to be *plausible* if *(1)* its topology is identical (RF=0) to the best-known ML tree or *(2)* its log-likelihood score is not significantly worse than the log-likelihood score of the best ML tree according to the AU test (*p <* 0.05). The “best-known ML tree” is the tree with the best likelihood among all inferred trees via all tree search methods, plus the true tree for simulated datasets. Before comparing likelihoods and conducting the AU test, PhyloSmew re-optimizes the branch lengths and model parameters for all inferred trees using the RAxML-NG `-evaluate` command. This is done to eliminate potential deviations among tools w.r.t. how they optimize and compute log-likelihoods (optimization *ϵ*, treatment of gaps etc.).

We evaluated the tree inference accuracy and speed of RAxML-NG 2.0 (*default* and *fast*) in comparison with the previous versions (RAxML-NG 1.1, RAxML-NG 1.2) as well as with two competing tools, IQTree (*default* and *fast*) and VeryFastTree (Table 3). We decided to benchmark Very-FastTree and not the older, more widely used FastTree2. because both tools share the same tree search heuristic, but VeryFastTree implements a more efficient parallelization scheme that can utilize more than 4 CPU cores.

**Table 3.** ML tree inference tools used for benchmarking.

| Abbr. | Tool | Version | Mode | Reference |
|---|---|---|---|---|
| RX1.1 | RAxML-NG | v1.1 | default | (1) |
| RX1.2 | RAxML-NG | v1.2 | default | (13) |
| RX2-DEF | RAxML-NG | v2.0 | default | (this paper) |
| RX2-FAST | RAxML-NG | v2.0 | -fast | (this paper) |
| IQ3-DEF | IQTree | v3.0.1 | default | (2) |
| IQ3-FAST | IQTree | v3.0.1 | -fast | (2) |
| VFT | VeryFastTree | v4.0.5 | -nosupport -double-precision | (6) |

## Results

### Tree search

In Figure 2, we show inference accuracy (percentage of plausible trees, RF distances) and speed (accumulated runtime, speedups) for each tool. In Figure S6 and Figure S7, we display the same metrics, but grouped by dataset type (empirical DNA, simulated DNA, empirical AA, simulated AA).

**Fig. 2.**
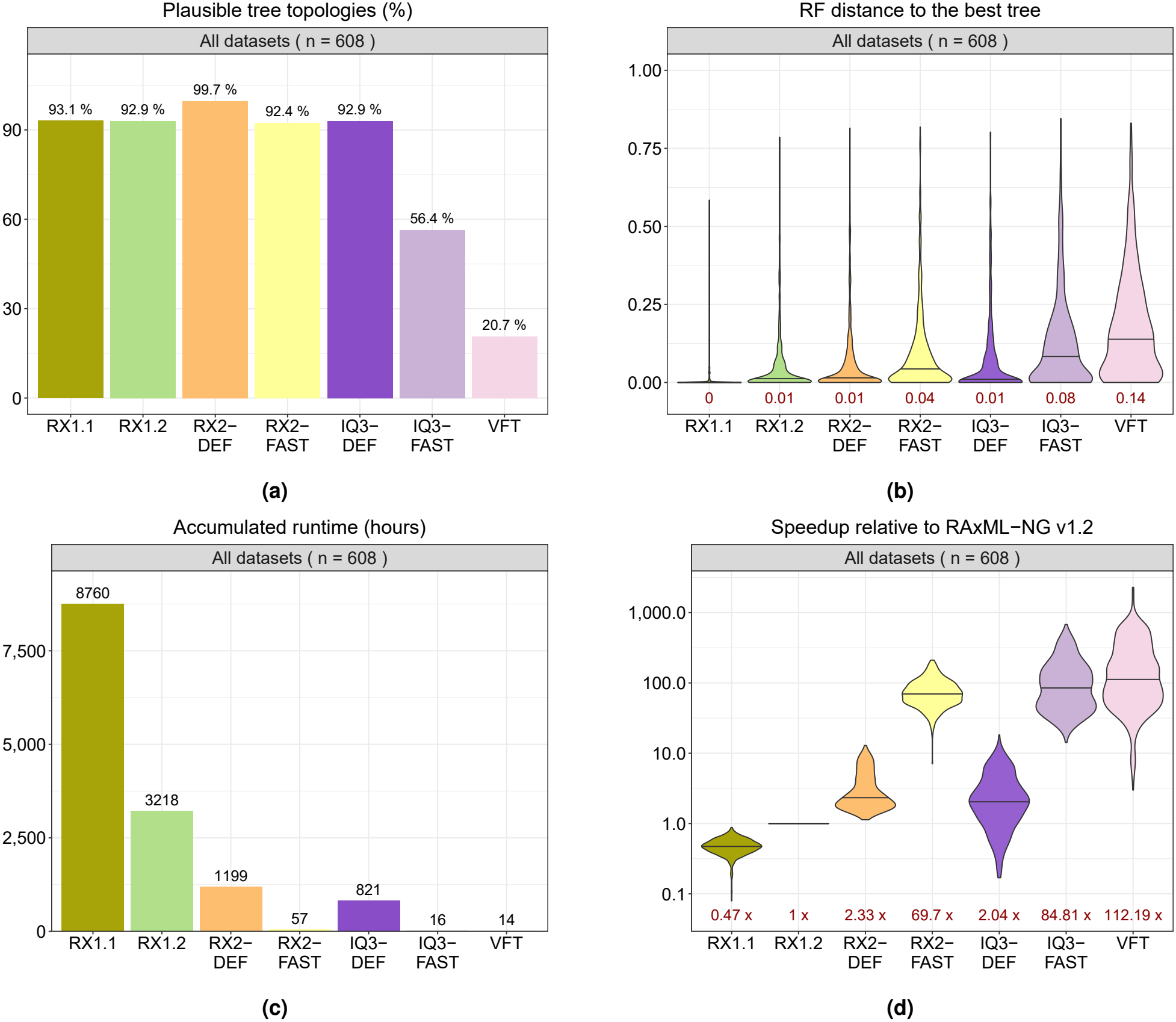
Tree search benchmark results: *(a)* Percentage of plausible trees, *(b)* Normalized RF distance to the best ML tree, *(c)* Accumulated runtime, *(d)* Speedup relative to RAxML-NG 1.2. Median values are shown in dark red.

In default mode (RX2-DEF), RAxML-NG 2.0 shows the highest accuracy both overall (99.7% plausible topologies) as well as for all four individual dataset types (98 − 100%), followed by RX1.1 with 93.2% (86 − 100%), RX1.2 with 92.9% (85 − 100%), IQ3-DEF with 92.9% (82 − 98%), and RX2-FAST with 92.4% (92 − 93%). IQ3-FAST and VeryFastTree yielded substantially lower accuracies of 56.4% (40 − 85%) and 20.7% (8 − 35%), respectively. All three fast methods (RX2-FAST, IQ3-FAST, and VFT) also exhibit higher RF distances to the best-known ML tree than their more thorough counterparts (median nRF: 0.04 − 0.14 vs. 0.00 − 0.01, see Figure 2b). It is worth noting that this pronounced difference in RF distances to the reference tree among thorough versus fast search methods can not be observed on simulated datasets (Figure S8) when the true tree is being used as a reference for computing RF distances. This phenomenon has been previously reported (31) and requires further investigation.

We use RX1.2, the latest production version of RAxML-NG, as a baseline for our runtime benchmark. RX1.1 is ≈ 2× slower, confirming our previous results from (13). RX2-DEF shows a median speedup of 2.33× (range: 1.13 − 12.88×), and IQ3-DEF a median speedup of 2.04× (range: 0.17 − 18.22×). Regarding the fast heuristics, VFT achieved the highest speedups (median: 112.19×, range: 2.99 − 2294.70×), followed by IQ3-FAST (median: 84.81×, range: 14.23 − 677.73×), and RX2-FAST (median: 69.70×, range: 7.11 − 210.97×).

### Model selection

We evaluate the performance and accuracy of our model selection tool MOOSE against ModelFinder (**?**), which forms part of IQTree (2).

To this end, we reuse the empirical datasets from the preceding tree search benchmarks. We use the default parameters of both tools with a fixed number of 8 threads. Additionally, for DNA datasets, we enable ML-optimized base frequencies for ModelFinder instead of using the default empirical frequencies that are derived by counting nucleotides in the input alignment. ModelFinder and MOOSE conduct model selection on distinct tree topologies, which renders the BIC (Bayesian Information Criterion) scores (37) reported by both tools incomparable with each other. Hence, we use IQTree to re-evaluate the models selected by both ModelFinder and MOOSE, on the fixed tree topology that ModelFinder used for model selection, and thereby obtain comparable BIC scores, Figure 3a shows the number and percentage of datasets where (1) both MOOSE and ModelFinder selected the same model, or (2) either tool selected a model with a lower (better) BIC score. If the difference in the BIC score exceeds 10, we consider that the model with the lower BIC score constitutes a significantly better fit.

**Fig. 3.**
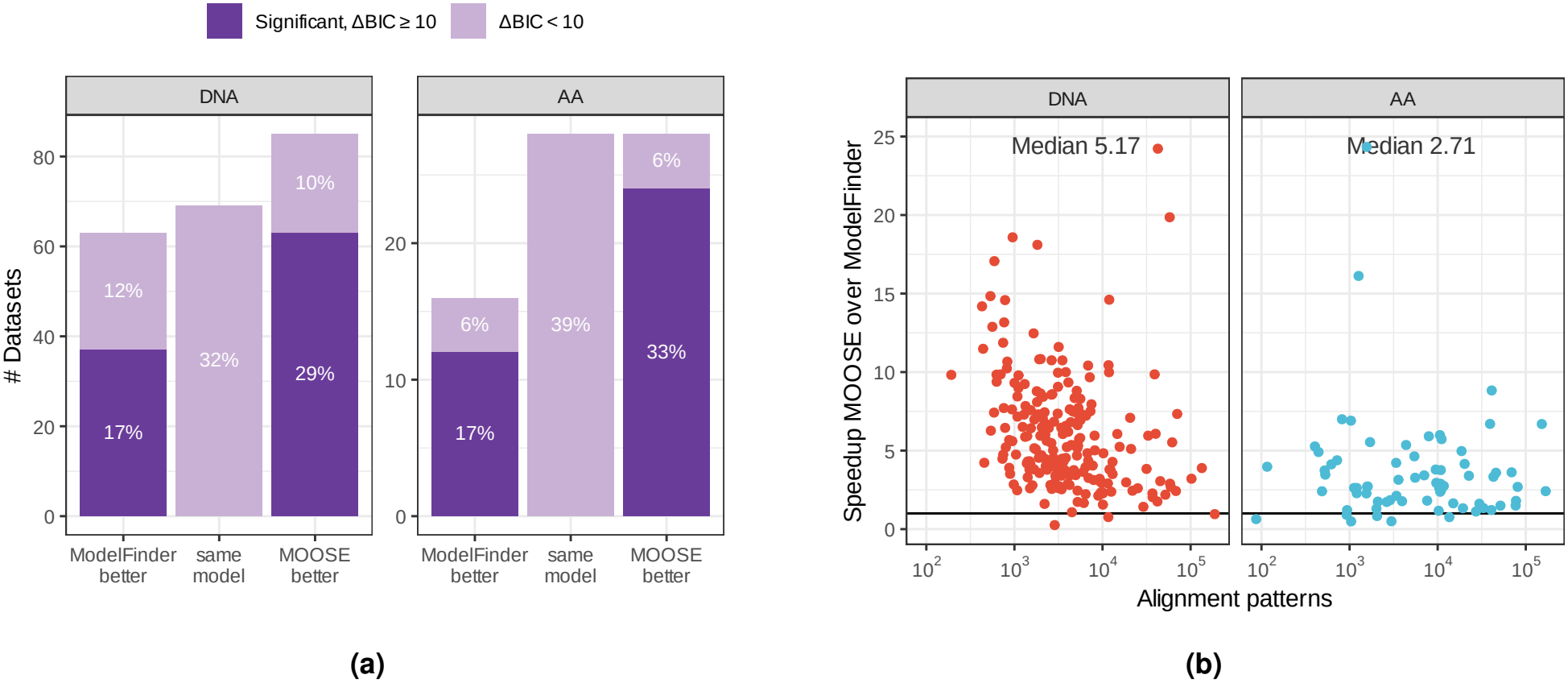
Speed and accuracy of MOOSE. ***(a)*** BIC score comparison between the model selected by ModelFinder and MOOSE after reevaluation with IQTree. ***(b)*** Speedup of MOOSE over ModelFinder over increasing alignment size.

MOOSE found a better fit model more frequently than ModelFinder (DNA: 29% vs. 17%, AA: 33% vs. 15%), although for the majority of datasets (DNA: 54%, AA: 52%) the differences in model fit were not significant. MOOSE was also substantially faster than ModelFinder with a median speedup of 5.4× on DNA and 2.8× on AA datasets.

### Branch support

We evaluated four branch support metrics: the standard Felsenstein bootstrap (FBP), the rapid bootstrap (RBS), the Educated Bootstrap Guesser (EBG-Light variant, henceforth: EBG), and the UltraFast bootstrap as implemented in IQTree3 (UFBoot). Due to excessive computational cost of the standard bootstrapping, we used a subset of 90 simulated datasets from the tree search benchmark (3 out of 10 replicates for each dataset size).

While FBP, RBS, and EBG can estimate branch supports on a pre-existing, fixed tree topology, UFBoot supports can only be computed during an IQTree3 tree search. Therefore, and to also better reflect typical usage scenarios, in our benchmark runs, we perform both tree search *and* branch support estimation. More specifically, we use the default RAxML-NG 2.0 search heuristic (RX2-DEF) in conjunction with computing FBP and RBS supports, the fast RAxMLNG 2.0 search heuristic (RX2-FAST) in combination with EBG, and the default IQTree3 search heuristic (IQ3-DEF) in combination with UFBoot. We do not include IQ3-FAST and VeryFastTree due to their inferior accuracy in our preceding tree search tests (see above).

We compared the branch support values inferred by each method with the probability of observing the respective branch in the true tree. The results are shown in Figure 4a. Overall, EBG support values are closest to the true branch probabilities for DNA datasets, but underestimate them for AA datasets. However, there is a substantial variation w.r.t. phylogentic difficulty and dataset size (see Supplementary results, Figures S9-S11). RBS and FBP yield very similar results, and both methods tend to overestimate the true branch probabilities. UFBoot exhibits the most pronounced overestimation of branch support. This is consistent with previous studies (19, 38, 39).

**Fig. 4.**
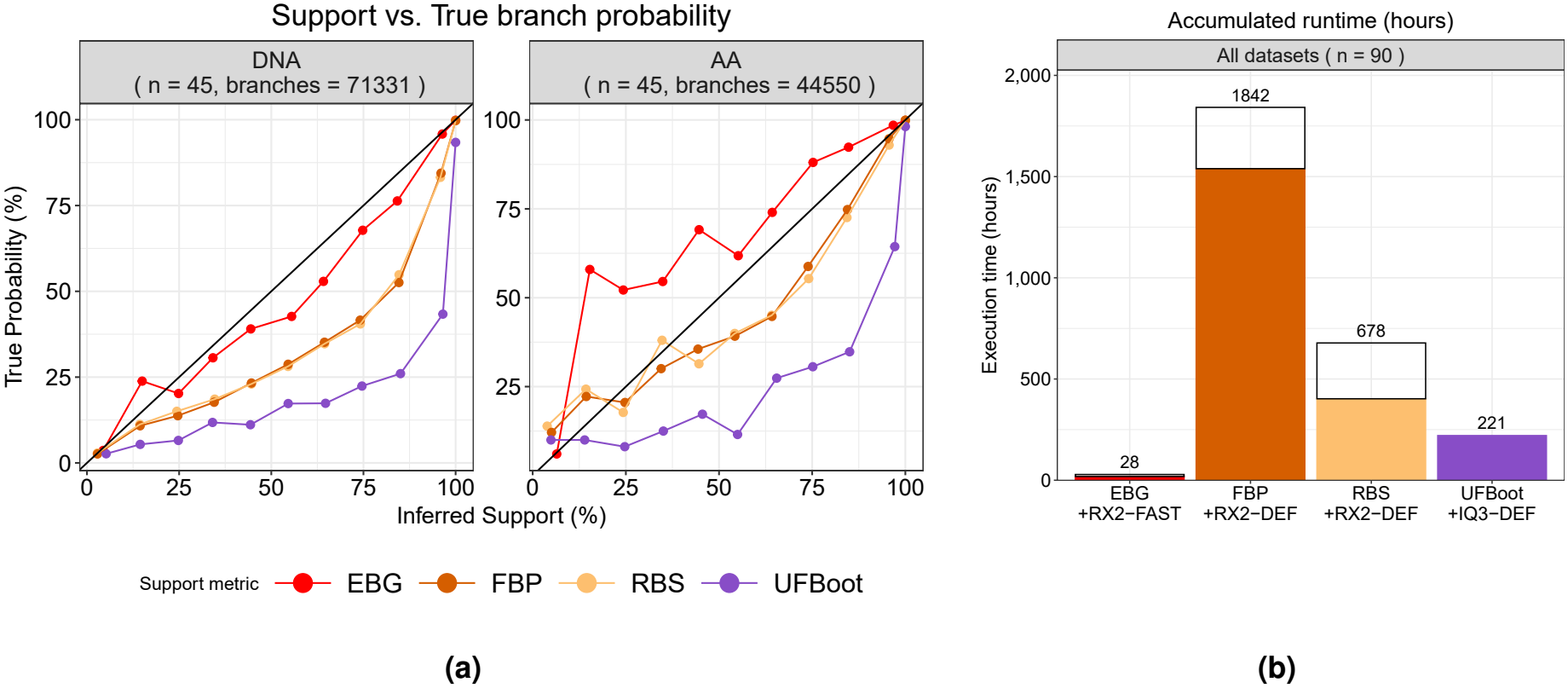
Speed and accuracy of fast branch support metrics. Evaluated on a subsample (*n* = 90) of the simulated DNA and AA datasets. ***(a)*** Estimated mean branch support vs. the actual proportion of branches in the true tree in bins of size 10. ***(b)*** Accumulated runtime for tree search (empty bars) and bootstrap support estimation (filled bars); UFBoot runtime cannot be separated from the tree search.

EBG in conjunction with RX2-FAST completed the bench-mark in about 28 hours which is ≈ 8× faster than IQTree3 with UFBoot, ≈ 24× faster than RBS+RX2-DEF, and ≈ 65× faster than FBP+RX2-DEF (Figure 4b). If we exclude the tree search phase and consider only branch support inference time, RBS is ≈ 3.8× faster than FBP and EBG is ≈ 85× faster than FBP.

## Discussion

Our evaluation shows that the default adaptive heuristic of RAxML-NG 2.0 matches and partially even exceeds the accuracy of RAxML-NG 1.2, while being 2 − 4× faster. The inference accuracy of the *fast* RAxML-NG 2.0 heuristic is only slightly inferior to RAxML-NG 1.2 and IQTree 3.0 (default mode), but requires an order of magnitude less runtime (20 − 30× speedup). Hence, we recommend RAxMLNG 2.0 *fast* as a more accurate alternative to VeryFastTree and IQTree *fast*, whenever a somewhat (3 − 5×) higher runtime compared to these methods is acceptable.

In our branch support benchmark, we observe that support estimation across all methods increases with a growing number of alignment sites (Figure S11). With 500 AA sites, FBP and RBS yield near-perfect estimates of true branch probabilities (curves are close to the diagonal), but with 1k to 8k AA sites, they show an increasingly severe support overestimation. EBG has the highest accuracy with 4k/8k AA sites, but underestimates support on smaller alignments. A similar trend is observed on DNA data (Figure S10): all methods overestimate support when the number of sites is sufficiently high (≥ 16k). UFBoot is least affected by dataset size and phylogenetic difficulty, but it also exhibits the lowest accuracy (consistent, pronounced overestimation).

In our model selection benchmark with 8 threads, MOOSE exhibits a considerable speedup over IQTree’s ModelFinder. Presumably, the coarse-grained parallelization across candidate models, an approach also adopted by ModelTest-NG, boosts the performance of MOOSE on datasets where the number of alignment sites does not suffice to efficiently utilize all CPU cores.

Phylogenomic studies have become commonplace, and there are three ways to use RAxML-NG for such genome-wide datasets. First, a concatenated alignment of all orthologous regions (“supermatrix”) can be analyzed with a partitioned model. RAxML-NG is highly efficient and scalable for concatenated matrices, as has been demonstrated in our previous work (1) and in a recent empirical study with a supermatrix comprising 63,430 genomic regions and ≈60 million sites (9). Alternatively, RAxML-NG can be used to infer gene trees, which are subsequently combined to obtain a species tree using methods such as ASTRAL (40), Species-Rax (41) or AleRax (42). The latter two, as well as our NetRAX tool (43) for hybridization-aware network inference, have a shared codebase with RAxML-NG. Hence, the new features described here, such as difficulty prediction or integrated model selection, can potentially also be adopted for these tools. Finally, branch support metrics such as Tree concordance factor (TCF) and Internode certainty (IC) allow to quantify the conflicting signal among genomic regions. Additional approaches, e.g., sliding-window bootstrapping (44), can be easily implemented via an external script.

In conclusion, our analysis shows that even well-established, decade-old heuristics still exhibit substantial optimization potential. Even higher performance gains can be attained when using classical machine-learning models that rely upon carefully engineered feature sets. Recently proposed tree inference methods based on deep learning (45, 46) show some promise, but are still too memory-intensive to scale beyond moderate dataset sizes (≈ 100 taxa). These limitations will likely be addressed and alleviated by future hardware and software improvements. Mechanistic models such as parsimony and continuous time Markov Chain substitution models, still have the conceptual advantage of explainability, but it remains to be seen which methods and tools will be adopted by practitioners in the years to come.

## Funding

This work was supported by the Klaus Tschira Foundation and by the European Union (EU) under Grant Agreement No. 101087081 (Comp-Biodiv-GR).

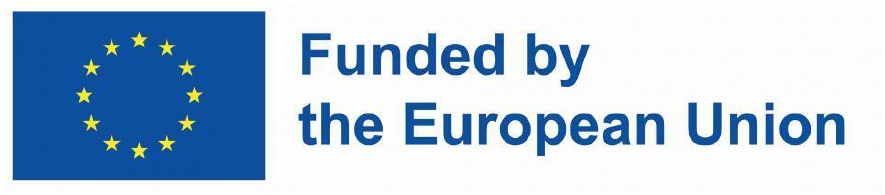

## Acknowledgment

We thank Johannes Hengstler and Luise Häuser for their helpful suggestions and for testing preliminary versions of the code.

## Data availability

All the original data and scripts necessary to reproduce the analyses reported in this study can be accessed through the Dryad link: https://doi.org/10.5061/dryad.k3j9kd5rw.

## Supplementary methods

### Tree search heuristics

Note: The description of tree search heuristics in this section is adapted from (48) and (47). The basic building block of RAxML-NG search heuristics is a so-called SPR round: all possible subtrees are subsequently removed from the currently best tree *T*_*b*_, re-insertions into all neighboring branches up to a specified rearrangement distance are performed, and corresponding induced topologies 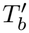 are scored. There are two scoring methods: in the *FAST SPR* mode, the log-likelihood of each induced topology with original branch length values is computed, whereas in the *SLOW SPR* mode three branches adjacent to the insertion point are additionally optimized using the Newton-Raphson method. If any of the induced topologies 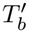 yields a higher log-likelihood score than the currently best tree, the corresponding SPR move is immediately applied, and we continue the optimization process with the new best tree 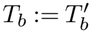.

### Default adaptive heuristic

The default search heuristic of RAxML-NG 2.0 is illustrated in Figure S2 and comprises the following steps:

1. Generate parsimony trees (default: 24) using randomized step-wise addition. These trees are required to compute Pythia features (see below), but will also be recycled as starting trees for ML search.
2. Use Pythia (15) to predict the phylogenetic difficulty *D* of the input MSA.
3. Determine the number of random and parsimony starting trees as a function of the difficulty *D*. For both random and parsimony starting trees, this function is a bell-shaped curve with a maximum value of 10 (parsimony) or 5 (random) at a difficulty score of 0.5 (Figure S1a).
4. Determine the adaptive SPR regrafting radius r_adaptive_ as a function of the difficulty score. This parameter starts from a value of 5 for a difficulty score of 0.0 and increases linearly to a value of 10 at difficulty 0.4. It then remains constant up to difficulty 0.6, after which it decreases linearly back to a value of 5 at a difficulty score of 1.0. The r_adaptive_ parameter determines the thoroughness of each independent tree inference and is multiplied by different factors in the two distinct phases of the tree search (see below).
5. For every starting tree, repeat steps 6-11.
6. Initial branch length optimization (BLO) with log-likelihood improvement threshold of *ε* := 10.
7. Initial model parameter optimization (MPO) with log-likelihood improvement threshold of *ε* := 10.
8. **FAST SPR phase**: Perform a sequence of *fast* SPR rounds with minimum and maximum SPR regrafting radii set to r_min_ := 1 and r_max_ := min(3 · r_adaptive_, 25), respectively. The sequence of FAST SPR rounds continues as long as the log-likelihood improvement ΔL := L^′^ − L between the tree T^′^ after the SPR round and the tree T before SPR round is significant, that is, it exceeds the threshold value *ε*. The default heuristic uses a fixed threshold of *ε* := 10, the same as in RAxML-NG 1.2. Alternatively, *ε* can be dynamically computed using a statistical test (see `--stop-rule` option).
9. Intermediate MPO with *ε* := 3.
10. **SLOW SPR phase**: Perform a sequence of *fast* SPR rounds. This phase is divided into two stages. In the first stage (Stage I), the SPR round is invoked with r_min_ := 1 and r_max_ := r_slow_ := min(r_adaptive_, 7), respectively. The tree T (with log-likelihood L) prior to Stage I is stored. After the SLOW SPR round, the resulting tree T^′^ (with log-likelihood L^′^) is also stored. The two trees T and T^′^ are then compared in an analogous way as in the FAST SPR rounds (i.e., either using the default threshold *ε* := 10, or a user-specified statistical test). If the improvement is significant, the algorithm returns to the state preceding Stage I, otherwise it proceeds to Stage II. In Stage II, the SLOW SPR round is invoked with r_min_ := r_max_ + 1 and r_max_ := 2 · r_max_, respectively. Only the resulting tree after the SPR round is stored as T^′^, while the reference tree T remains the tree prior to Stage I. The two trees T and T^′^ are again compared. If the log-likelihood improvement is significant, the algorithm returns to the state preceding Stage I. Otherwise, the algorithm exits the sequence of SLOW SPR rounds. In other words, the sequence of SLOW SPR rounds only terminates when two consecutive SPR rounds (the first with r_min_ := 1 and r_max_ := min(r_adaptive_, 7), and the second with r_min_ := r_slow_ + 1 and r_max_ := 2 · r_slow_, respectively) yield non-significant log-likelihood improvement.
11. Final MPO with *ε* := 0.1.
12. Select the tree with the highest log-likelihood score as the best-known ML tree.

In both FAST and SLOW SPR rounds, the default heuristic maintains a list of the *X* := 20 best-scoring topologies encountered during the search, which it re-evaluates with full BLO after the SPR round. This parameter setting is identical to that used in RAxML-NG 1.2. Moreover, in both FAST and SLOW SPR rounds, the subtree cutoff feature is enabled, with an initial threshold value of 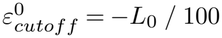. This initial threshold is higher than in RAxML-NG 1.2, where the corresponding value was −*L*_0_ */* 1, 000. Practically, this increase results in a larger number of SPR topologies being evaluated during the initial invocation(s) of both the FAST and SLOW SPR rounds. RAxML-NG 2.0 resets the subtree cutoff threshold to its initial value after each MPO round, that is, before entering the FAST and SLOW SPR phases.

### Fast heuristic

The fast search heuristic of RAxML-NG 2.0 is illustrated in Figure S3 and comprises the following steps:

**Fig. S1.**
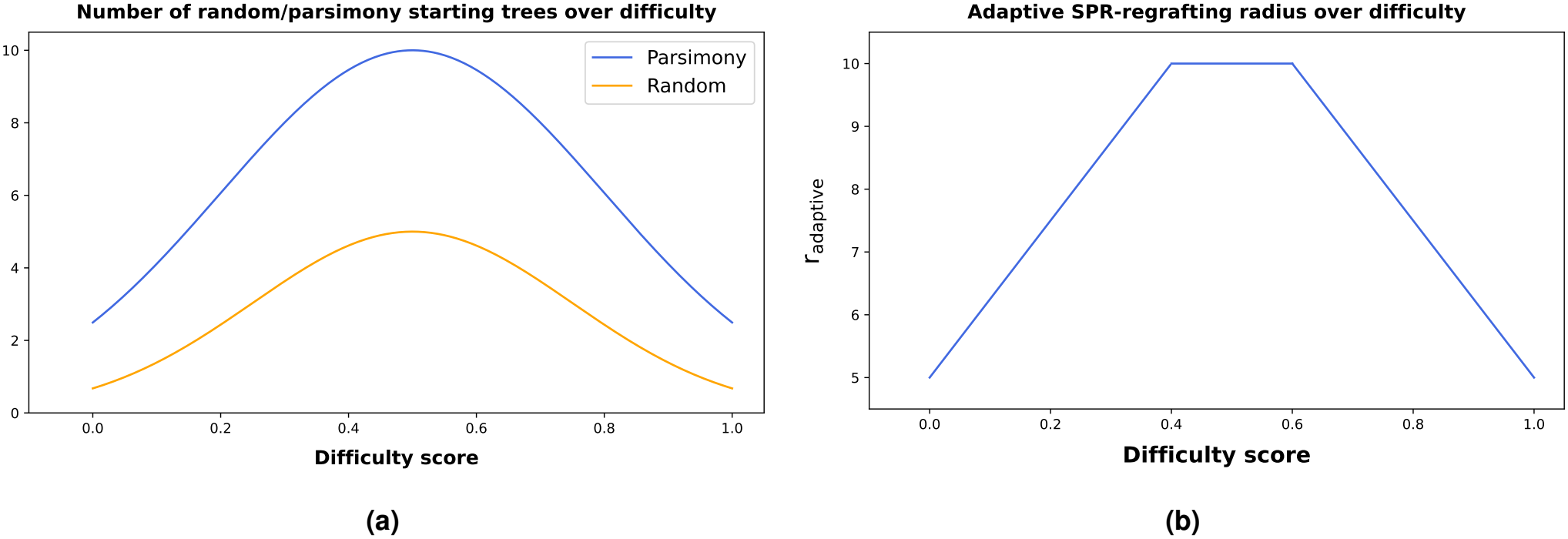
Adaptive number of random and parsimony starting trees *(a)* and SPR-regrafting radius *(b)* as a function of the Pythia difficulty score in the default tree search heuristic of RAxML-NG 2. Original figure from (47).

1. Initial branch length optimization (BLO) with log-likelihood improvement threshold of *ε* := 10.
2. Initial model parameter optimization (MPO) with log-likelihood improvement threshold of *ε* := 10.
3. **The FAST SPR phase** Perform a sequence of fast SPR rounds with a fixed radius of 10 (r_min_ := 1 and r_max_ := 10). By default, the log-likelihood improvement threshold *ε*_*KH*_ is calculated using the KH statistical test with multiple-testing correction (see (16) for details).
4. Intermediate MPO with *ε* := 3.
5. **The SLOW SPR phase** Perform a sequence of fast SPR rounds with a fixed radius of 10 (r_min_ := 1 and r_max_ := 10). By default, the log-likelihood improvement threshold *ε*_*KH*_ is calculated using the KH statistical test with multiple-testing correction.
6. Final MPO with *ε* := 0.1.

### Legacy search heuristics

It is possible to invoke the “classic” search heuristic of RAxML-NG 1.x via the `--opt-topology classic` option. However, this will not affect the number of starting trees. To fully reproduce the search behavior of RAxML-NG 1.1 and RAxML-NG 1.2, use `--extra compat-v11` and `--extra compat-v12`, respectively. It is possible to invoke the “classic” search heuristic of RAxML-NG 1.x via the `--opt-topology classic` option. However, this will not affect the number of starting trees. To fully reproduce the search behavior of RAxML-NG 1.1 and RAxML-NG 1.2, use `--extra compat-v11` and `--extra compat-v12`, respectively.

### EBG-Light

Educated Bootstrap Guesser predicts branch support values using the features extracted from the MSA and the respective (ML) reference tree. The original EBG implementation used 23 features for the prediction (19). In EBG-Light, we reduce this set to five most essential features (Table S1):

- **Parsimony Support (PS)** We generate 1000 parsimony trees by randomly re-shuffling sequences in the original MSA and conducting parsimony-based step-wise addition procedure. For every branch in the reference tree, we compute its support as the percentage of parsimony trees that contain this branch.
- **Parsimony Bootstrap Support (PBS)** We generate 200 replicate MSAs by resampling the columns of the original MSA with replacement. For each of the replicate MSAs, we conduct parsimony-based stepwise addition procedure to obtain the corresponding replicate tree. For every branch in the reference tree, we compute its support as the percentage of replicate trees that contain this branch.
- **Branch length** in the reference tree
- **Skewness PBS** The skewness of the PBS of all inner branches of the reference tree.
- **Mean RF distance PB** Average normalized Robinson-Foulds distance between parsimony bootstrap trees.

**Table S1.**
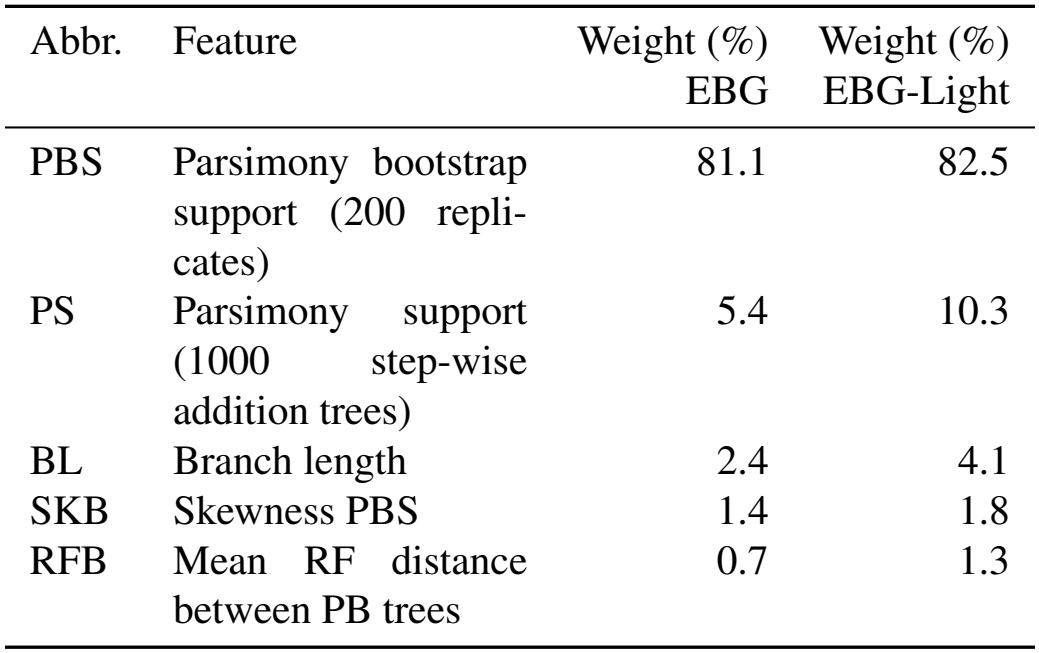
EBG-Light features.

| Abbr. | Feature | Weight (%)<br>EBG | Weight (%)<br>EBG-Light |
|---|---|---|---|
| PBS | Parsimony bootstrap support (200 repli-cates) | 81.1 | 82.5 |
| PS | Parsimony support (1000 step-wise addition trees) | 5.4 | 10.3 |
| BL | Branch length | 2.4 | 4.1 |
| SKB | Skewness PBS | 1.4 | 1.8 |
| RFB | Mean RF distance between PB trees | 0.7 | 1.3 |

**Fig. S2.**
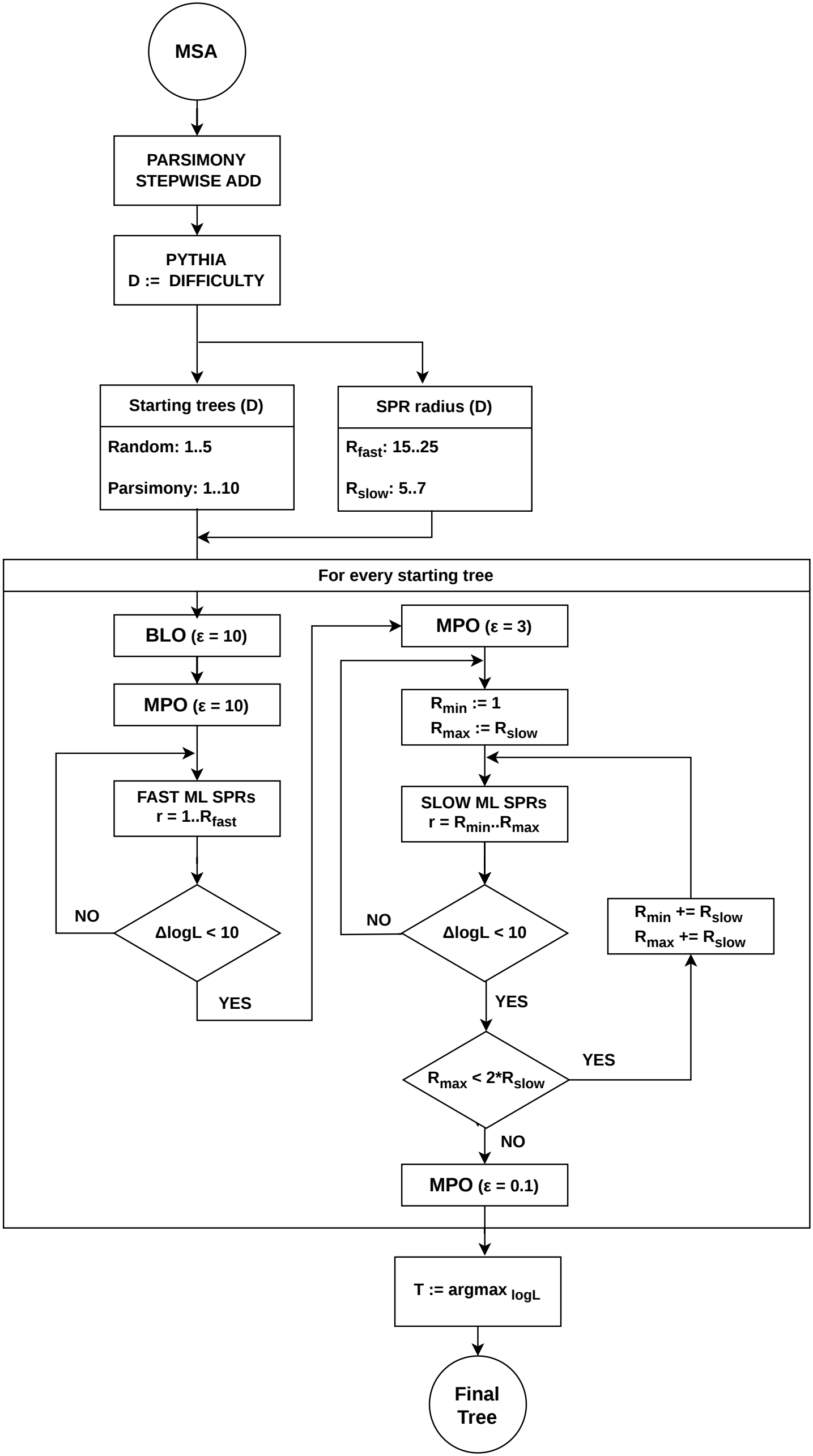
Default tree search heuristic in RAxML-NG 2.0. Abbreviations: BLO = Branch Length Optimization, MPO = Model Parameter Optimization, SPR = Subtree Pruning and Regrafting.

**Fig. S3.**
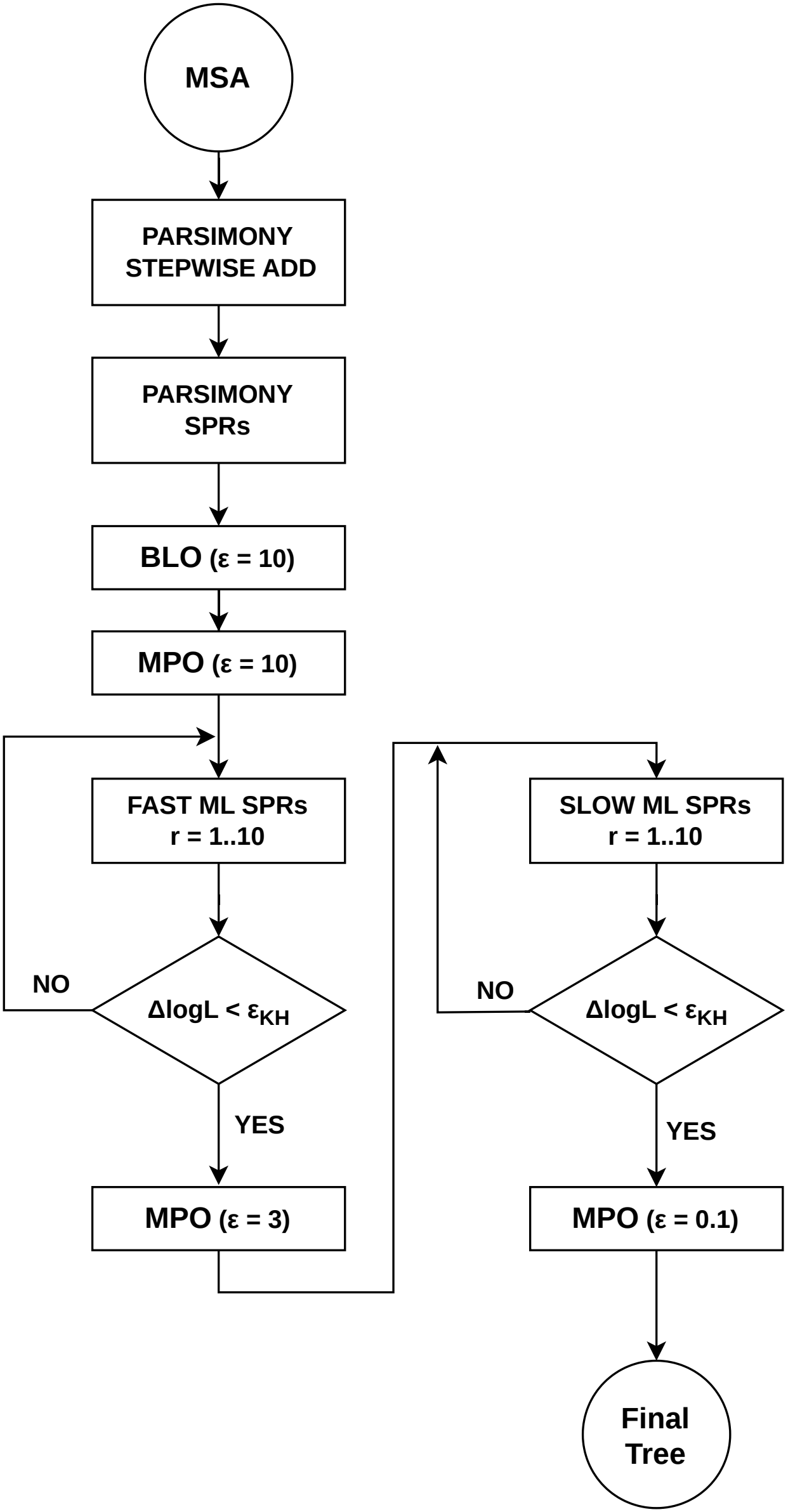
Fast tree search heuristic in RAxML-NG 2.0. Abbreviations: BLO = Branch Length Optimization, MPO = Model Parameter Optimization, SPR = Subtree Pruning and Regrafting.

### MOOSE

#### Heuristics

MOOSE follows the classical model selection approach which ranks models according to an information criterion (IC) such as BIC (37), and picks the model with the best IC score as the best-fit model. This process requires model optimization which becomes computationally expensive for large MSAs and/or for models with many free parameters. This is exacerbated by the high number of alternative substitution matrices and rate heterogeneity across sites (RHAS) models. Hence, MOOSE deploys heuristics to reduce the set of candidate models for evaluation. First, it evaluates all RHAS models on a single *reference* substitution matrix (`GTR` for DNA data, `LG` for AA data). Then, with the remaining substitution matrices, it will only test those RHAS models that were “promising” on the reference matrix (default: IC score difference of at most 10 to the best-observed score). Additionally, since FreeRate RHAS models have a variable number of categories, we start with the minimum (default: 2) and subsequently increase the number of categories until the IC score shows no further improvement or the maximum number (default: 10) is reached. This is illustrated in Figure S4a. The aforementioned heuristics are similar to those implemented by IQ-TREE’s ModelFinder (27), however, MOOSE features a more sophisticated parallelization strategy, as described below.

#### Evaluation Order & Parallelization

MOOSE supports concurrent evaluation of candidate models (coarse-grained parallelization), as well as distributing the computation of a single candidate model across multiple processor cores by splitting up the alignment sites (fine-grained parallelization), as illustrated in Figure S4b. As the heuristics described above introduce a soft dependency between candidate model evaluations (e.g. the reference substitution model should be evaluated before all others), all candidate models are ranked according to a priority and evaluated in that order. MOOSE applies the heuristics opportunistically, i.e. if the necessary results needed to determine whether a heuristic applies to a candidate model are not yet available, the model will be evaluated regardless, sometimes superfluously. To keep the number of superfluous model evaluations on partitioned datasets low, MOOSE reorders candidate models across partitions according to their priority, thereby increasing time between the evaluation reference models and the remaining evaluations. The priority depends on whether or not the substitution model is the reference model and on the rate category count being common, i.e. frequently chosen in the best-fit model: reference models with fewer than five categories have `HIGHEST` priority and with at least five categories `HIGH` priority, whereas the remaining models with fewer than five categories have `NORMAL` priority and with at least five categories `LOW` priority. Along with the number of alignment sites *N*, number of rate categories *C*, and state count *S*, the priority determines the assigned number of processor cores *p*^rounded^:

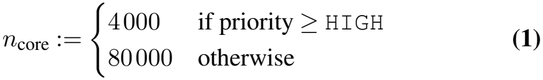

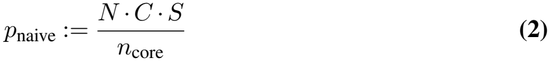

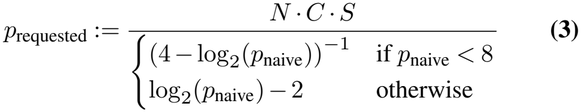

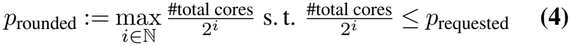

#### Model Parameter Optimization

To speed up the convergence of RHAS parameter optimization, MOOSE uses the parameter estimates of the reference model as a starting point for the optimization of the remaining substitution models. This includes the Gamma shape parameter *α*, FreeRate category rates and weights, and the proportion of invariant sites. For faster evaluation of FreeRate models in particular, RAxML-NG 2.0 implements the Expectation-Maximization (EM) algorithm introduced by Kalyaanamoorthy et al. (27) in addition to L-BFGS-B (49). In our experiments (data not shown), L-BFGS-B often obtains a better likelihood than EM, therefore it remains the default optimization algorithm for FreeRate models during tree search despite its higher runtime.

**Fig. S4.**
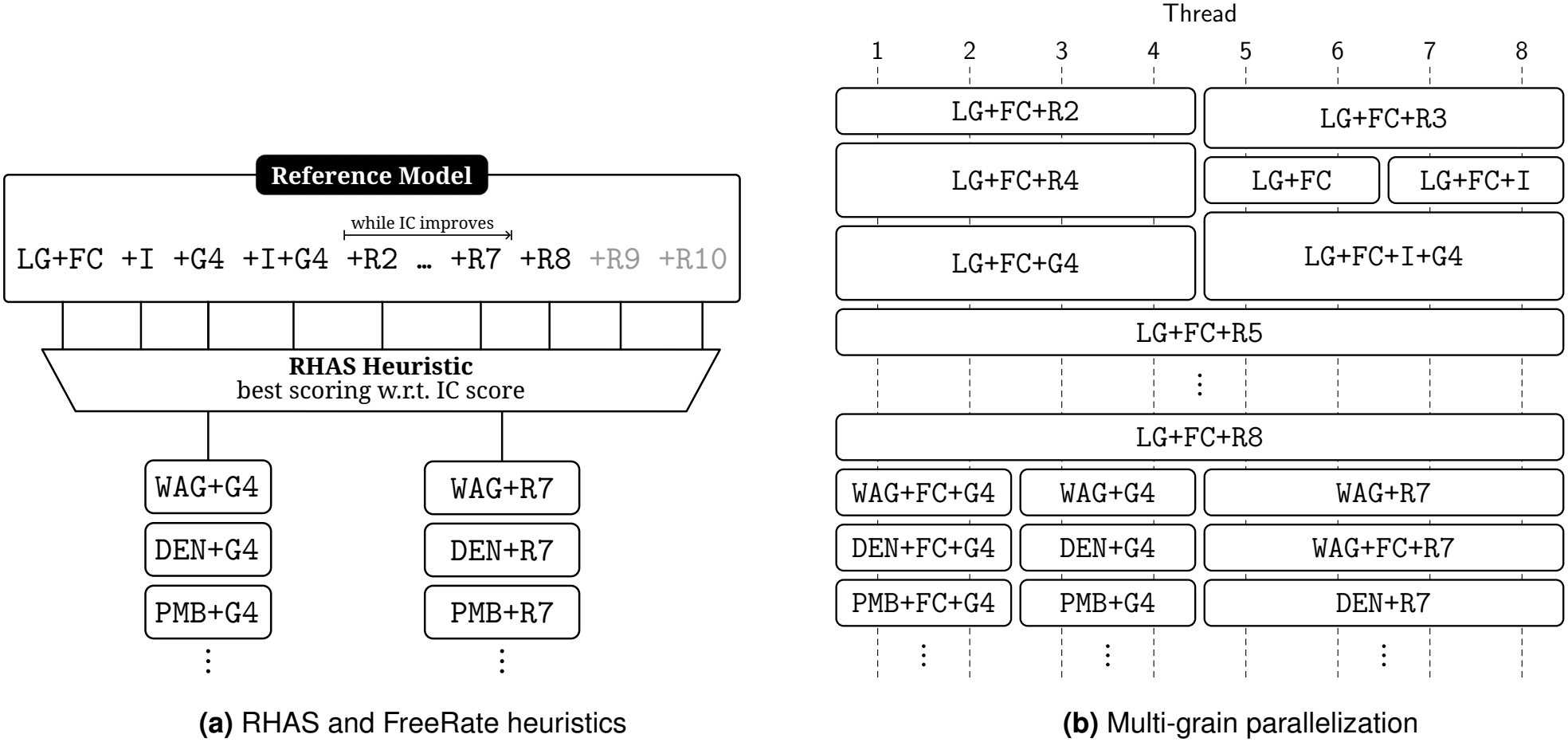
MOOSE candidate selection heuristics *(a)* and multi-grain parallelization *(b)* illustrated on a fictional protein MSA. For protein data, LG substitution matrix with empirical amino acid frequencies (`LG+FC`) is used as a reference model. All RHAS models (`+I`, `+G4`, `+I+G4`, `+R`) are evaluated on this reference model. For the FreeRate model, no significant IC score improvement has been observed with 8 categories (`+R8`), therefore `+R7` is considered the optimal number of FreeRate categories, and models with higher category counts (`+R9`, `+R10`) have been skipped (not evaluated). The difference in IC scores between `+R7` and `+G4` is below the significance threshold (default: 10), thus both RHAS models will be evaluated with the remaining protein substitution matrices (`WAG`, `DEN` etc.). Candidate models with higher number of categories are more expensive to evaluate, and hence they get assigned more threads: e.g., 4 threads for `WAG+R7` and 2 threads for `WAG+G4`. Furthermore, reference models get higher priority and therefore more resources: e.g., 8 threads for `LG+FC+R5` and 4 threads for `LG+FC+G4`.

### Tree Concordance Factor

Tree Concordance Factor (TCF) of branch *x* is defined as a proportion of gene trees that are *concordant* with *x* among all gene trees *decisive* for *x* (20). In other words, it is a proportion of gene trees that *do* contain a branch *x* among all gene trees that *could possibly* contain a branch *x*. If the reference (species) tree and all gene trees share the same taxon set, then every gene tree is decisive for every branch. In this case, TCF is equivalent to Gene Concordance Factor (gCF) as implemented in IQTree (22). However, in practice, certain genes can only be present in a subset of taxa, and hence respective gene trees are incomplete (contain fewer taxa than the species tree). In such cases, IQTree identifies a gene tree as *decisive* iif it contains at least one taxon from each of the four subtrees adjacent to *x* (see Figure S5). Under this definition, a gene tree that contains a branch incompatible with *x* can still be considered non-decisive and ignored, potentially yielding inflated support values. With TCF as implemented in RAxML-NG, we address this issue by re-defining decisiveness as follows: a gene tree is decisive for the branch *x* iif the corresponding induced split is non-trivial. In other words, a gene tree is decisive iif after pruning all taxa not present in the gene tree (induced split), each side contains at least 2 taxa (non-trivial).

### Genotype models

Character state encoding for the phased (GT16) and unphased (GT10) genotype models is shown in Table S2.

**Table S2.**
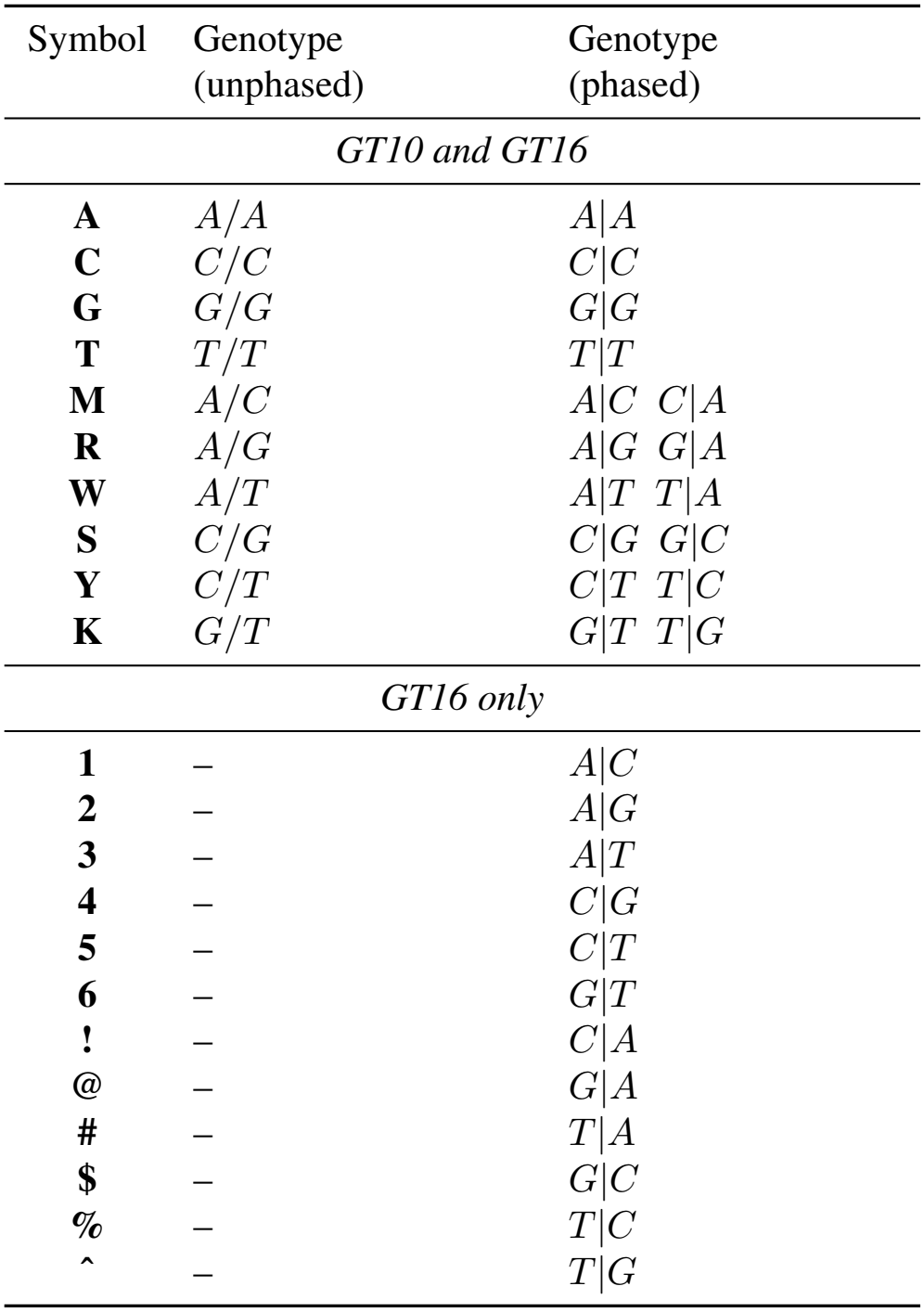
State encoding for genotype models.

| Symbol | Genotype<br>(unphased) | Genotype<br>(phased) |
|---|---|---|
| *GT10 and GT16* |  |  |
| **A** | *A/A* | *A\|A* |
| **C** | *C/C* | *C\|C* |
| **G** | *G/G* | *G\|G* |
| **T** | *T/T* | *T\|T* |
| **M** | *A/C* | *A\|C C\|A* |
| **R** | *A/G* | *A\|G G\|A* |
| **W** | *A/T* | *A\|T T\|A* |
| **S** | *C/G* | *C\|G G\|C* |
| **Y** | *C/T* | *C\|T T\|C* |
| **K** | *G/T* | *G\|T T\|G* |
| *GT16 only* |  |  |
| **1** | – | *A\|C* |
| **2** | – | *A\|G* |
| **3** | – | *A\|T* |
| **4** | – | *C\|G* |
| **5** | – | *C\|T* |
| **6** | – | *G\|T* |
| **!** | – | *C\|A* |
| **@** | – | *G\|A* |
| **#** | – | *T\|A* |
| **\$** | – | *G\|C* |
| **%** | – | *T\|C* |
| **^** | – | *T\|G* |

**Fig. S5.**
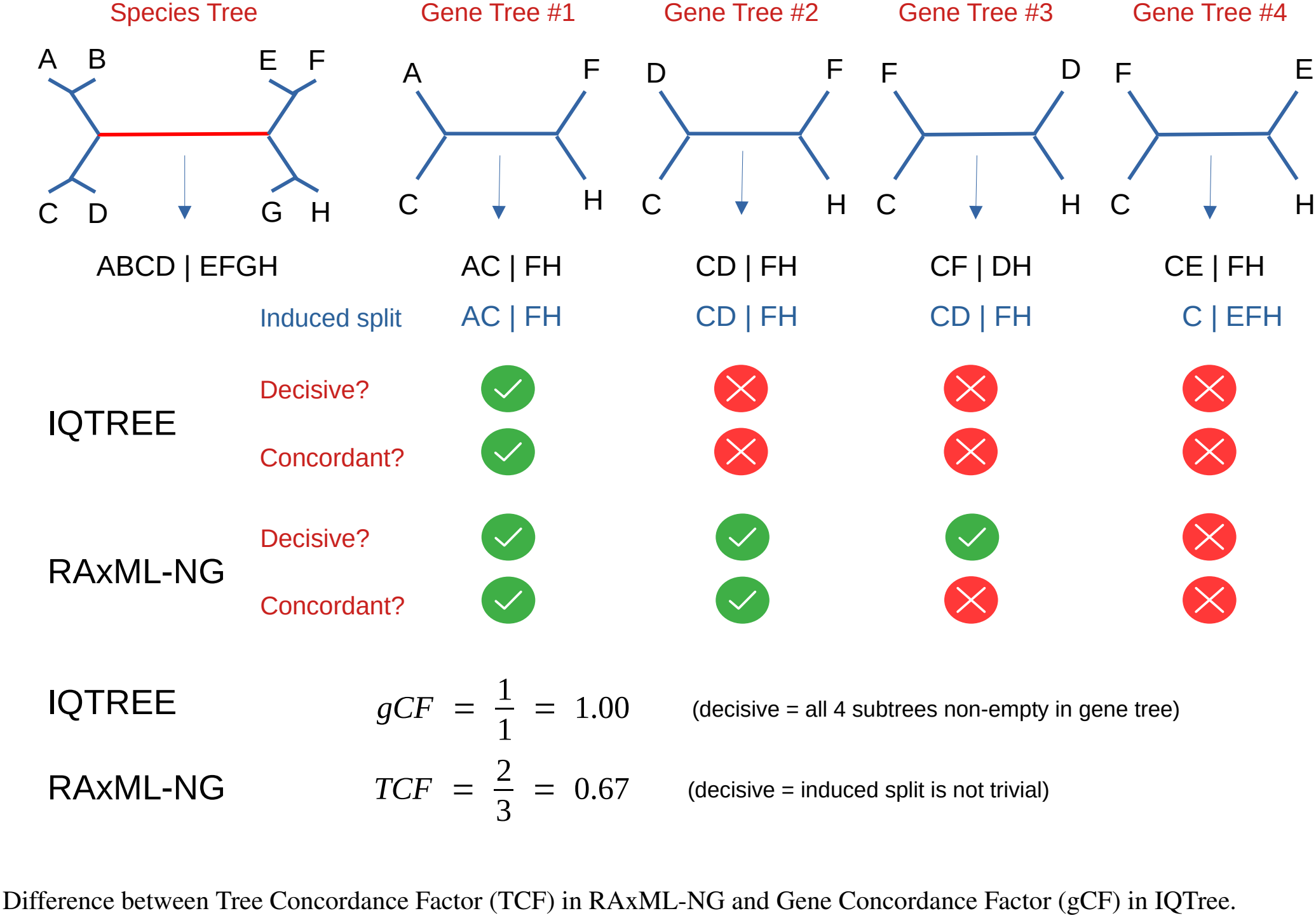
Difference between Tree Concordance Factor (TCF) in RAxML-NG and Gene Concordance Factor (gCF) in IQTree.

### Datasets

#### Empirical datasets

We used the following empirical datasets in our evaluation:

- 300 largest MSAs from TreeBase (30).
- 8 large protein families from InterPro (50) (see Table S3)
- 1 alignment of 2837 COI sequences from BOLD (51)

**Table S3.**
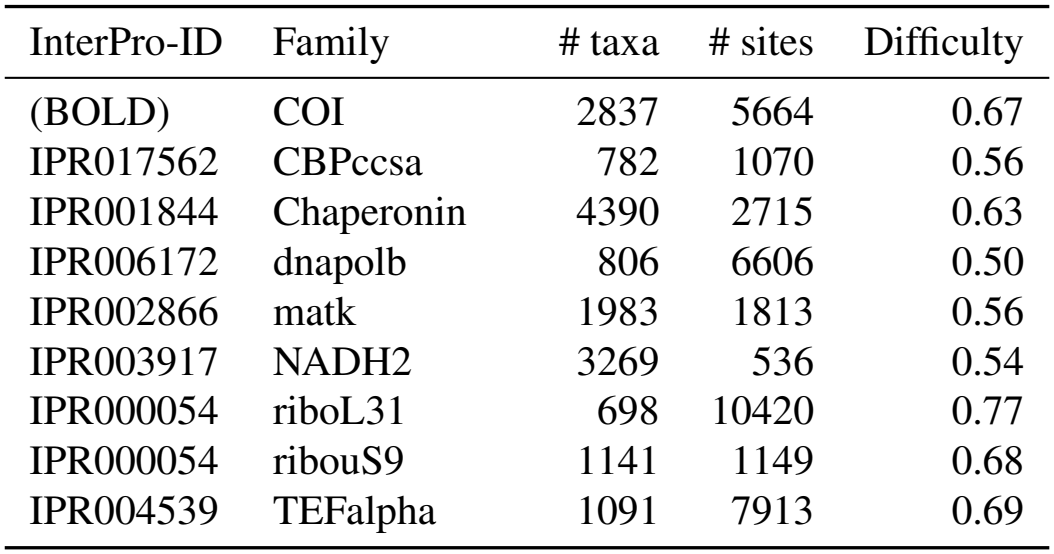
Large protein families used for benchmarking.

#### Simulated datasets

The concrete criteria used for the selection of reference datasets from RAxMLGrove for simulations were the following: For a target taxa number value *t*_*n*_, we selected datasets from RAxMLGrove with taxa number *t*_*rg*_ with the condition *t*_*n*_ ≤ *t*_*rg*_ ≤ `max`(*t*_*n*_ + 50, *t*_*n*_ × 1.05) for AA datasets, and *t*_*n*_ ≤ *t*_*rg*_ *<* `max`(*t*_*n*_ + 50, *t*_*n*_ × 2) for DNA datasets. The target values for AA taxon numbers were smaller than the target values for DNA datasets, such that it was easier to find datasets within the target range for AA, which is why we relaxed the formula for DNA datasets. From the candidates satisfying these conditions, we pulled 10 datasets at random with replacement. Then, we pruned taxa up to *t*_*n*_ uniformly at random.

#### Command lines

Specific command lines used for tree search, model selection, and branch support estimation are given in Table S4.

## Supplementary results

### Tree search

Figure S6. The accuracy and speed of the tree search in each dataset category are shown in Figure S6. RF distances to the the best ML tree and per-dataset speedups are shown in Figure S7. RF distances to the true tree on simulated datasets are shown in Figure S8.

### Branch support

Support metric accuracy binned by phylogenetic difficulty is shown in Figure S8. Support metric accuracy binned by the number of MSA sites is shown in Figure S10 (DNA) and Figure S11 (AA).

**Table S4.**
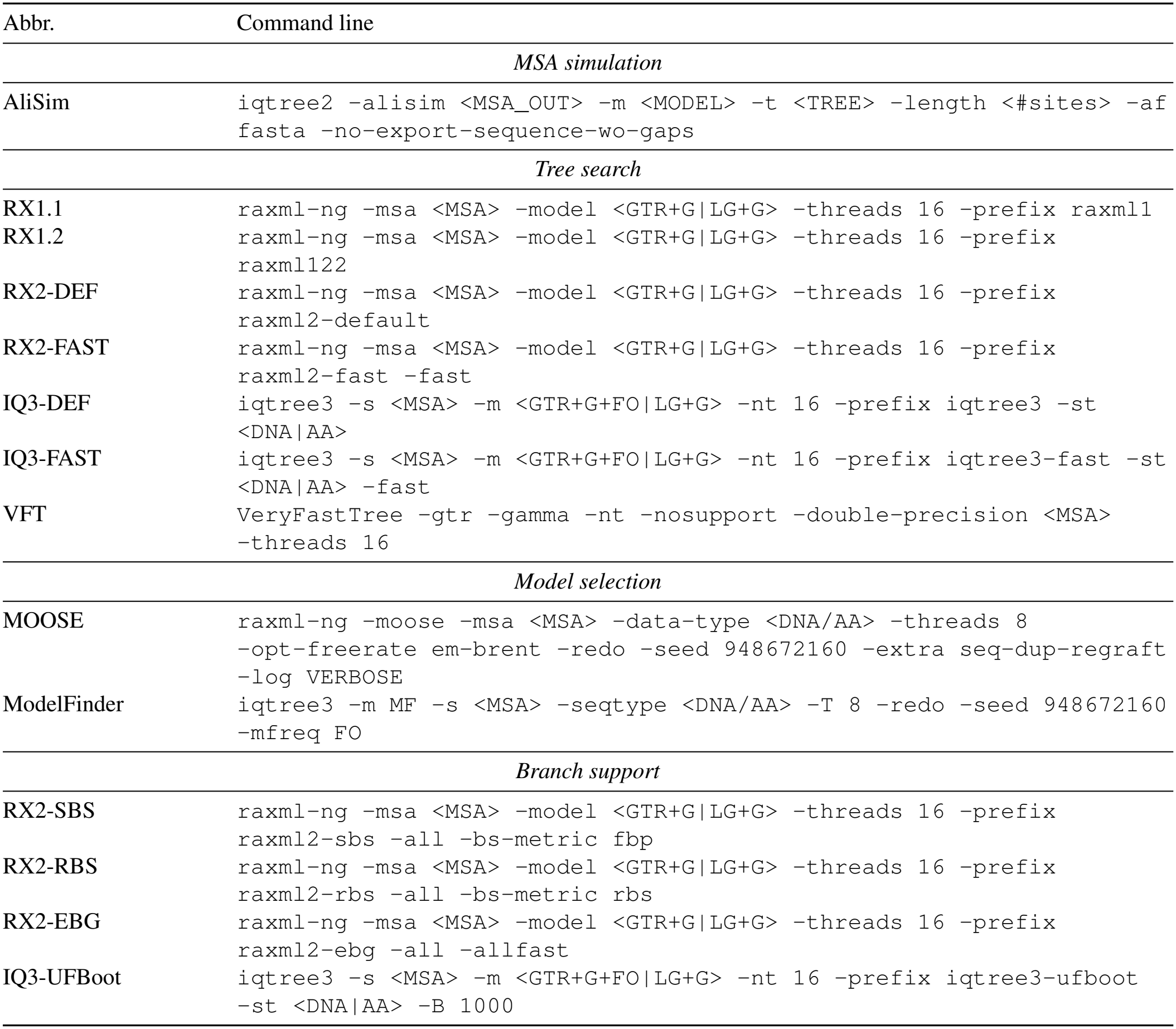
Command lines used for benchmarking.

| Abbr. | Command line |
|---|---|
| *MSA simulation* |  |
| AliSim | iqtree2 -alisim <MSA_OUT> -m <MODEL> -t <TREE> -length <#sites> -af fasta -no-export-sequence-wo-gaps |
| *Tree search* |  |
| RX1.1 | raxml-ng -msa <MSA> -model <GTR+G\|LG+G> -threads 16 -prefix raxml1 |
| RX1.2 | raxml-ng -msa <MSA> -model <GTR+G\|LG+G> -threads 16 -prefix raxml122 |
| RX2-DEF | raxml-ng -msa <MSA> -model <GTR+G\|LG+G> -threads 16 -prefix raxml2-default |
| RX2-FAST | raxml-ng -msa <MSA> -model <GTR+G\|LG+G> -threads 16 -prefix raxml2-fast -fast |
| IQ3-DEF | iqtree3 -s <MSA> -m <GTR+G+FO\|LG+G> -nt 16 -prefix iqtree3 -st <DNA\|AA> |
| IQ3-FAST | iqtree3 -s <MSA> -m <GTR+G+FO\|LG+G> -nt 16 -prefix iqtree3-fast -st <DNA\|AA> -fast |
| VFT | VeryFastTree -gtr -gamma -nt -nosupport -double-precision <MSA> -threads 16 |
| *Model selection* |  |
| MOOSE | raxml-ng -moose -msa <MSA> -data-type <DNA/AA> -threads 8 -opt-freerate em-brent -redo -seed 948672160 -extra seq-dup-regraft -log VERBOSE |
| ModelFinder | iqtree3 -m MF -s <MSA> -seqtype <DNA/AA> -T 8 -redo -seed 948672160 -mfreq FO |
| *Branch support* |  |
| RX2-SBS | raxml-ng -msa <MSA> -model <GTR+G\|LG+G> -threads 16 -prefix raxml2-sbs -all -bs-metric fbp |
| RX2-RBS | raxml-ng -msa <MSA> -model <GTR+G\|LG+G> -threads 16 -prefix raxml2-rbs -all -bs-metric rbs |
| RX2-EBG | raxml-ng -msa <MSA> -model <GTR+G\|LG+G> -threads 16 -prefix raxml2-ebg -all -allfast |
| IQ3-UFBoot | iqtree3 -s <MSA> -m <GTR+G+FO\|LG+G> -nt 16 -prefix iqtree3-ufboot -st <DNA\|AA> -B 1000 |

**Fig. S6.**
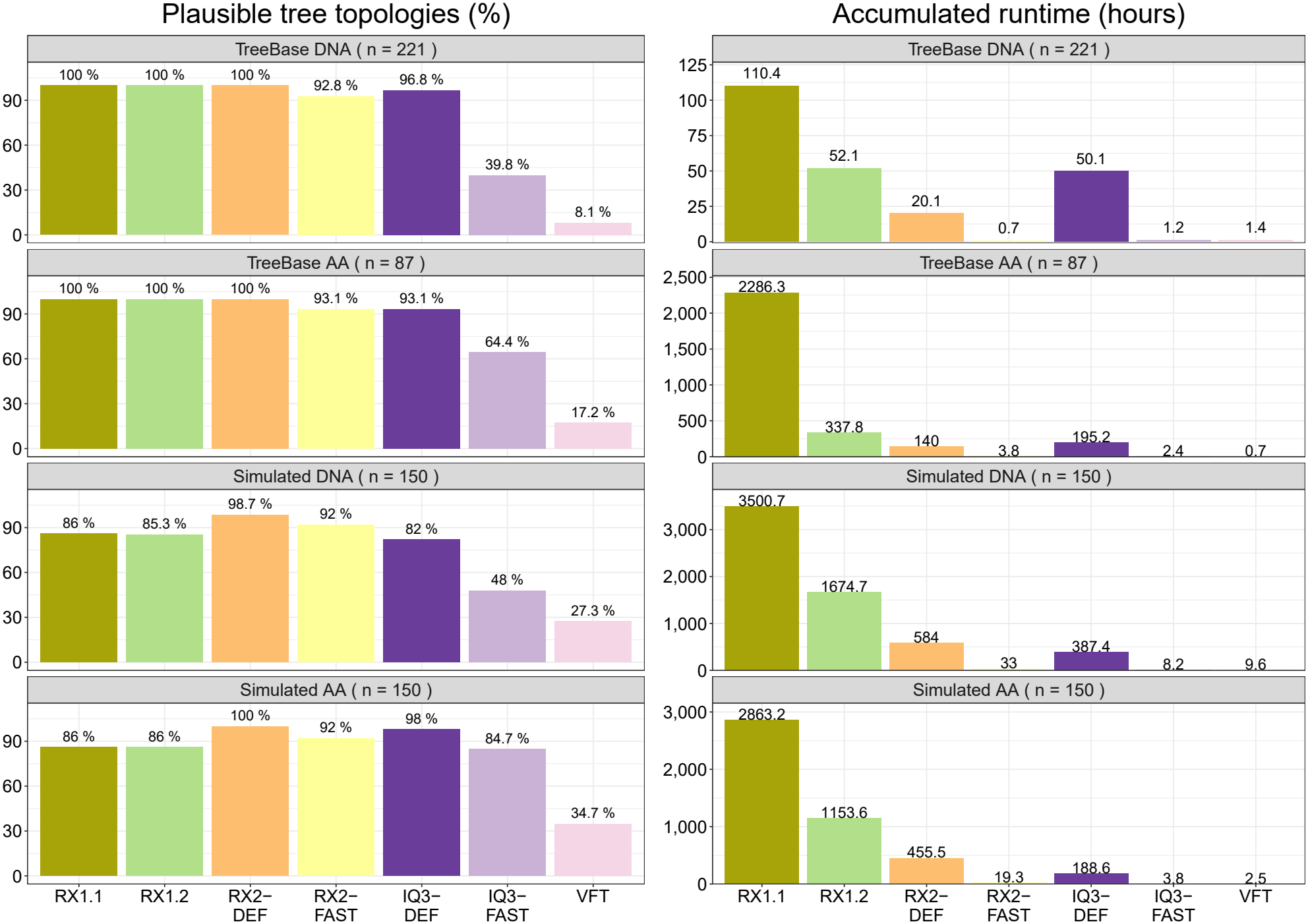
Tree search benchmark results. Percentage of plausible trees (left) and accumulated runtime (right).

**Fig. S7.**
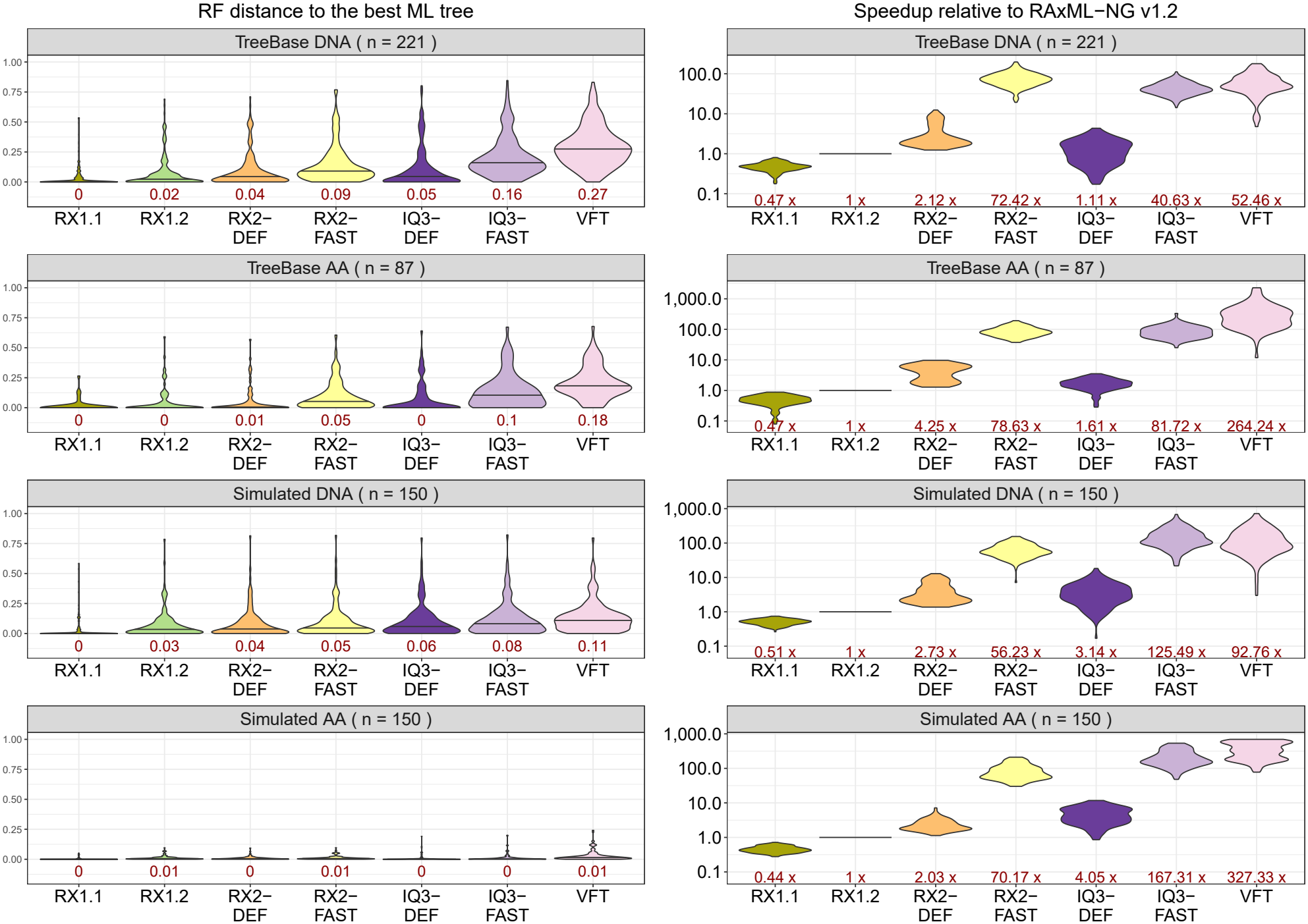
Tree search benchmark results. Normalized RF distance to the best ML tree (left) and speedup compared to RAxML-NG 1.2 (right). Median values shown in red.

**Fig. S8.**
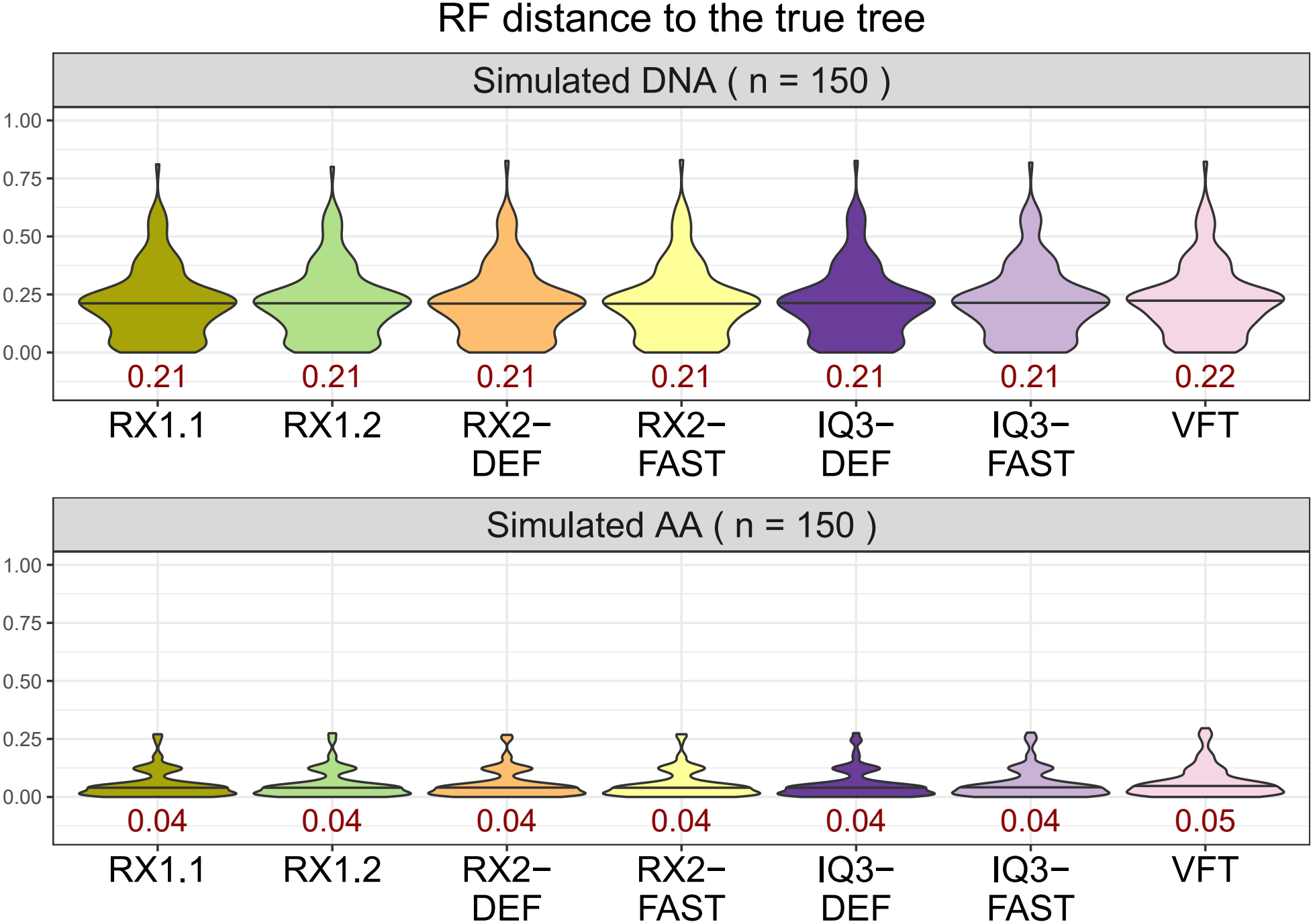
Normalized RF distance to the true tree on simulated datasets. Median values shown in red.

**Fig. S9.**
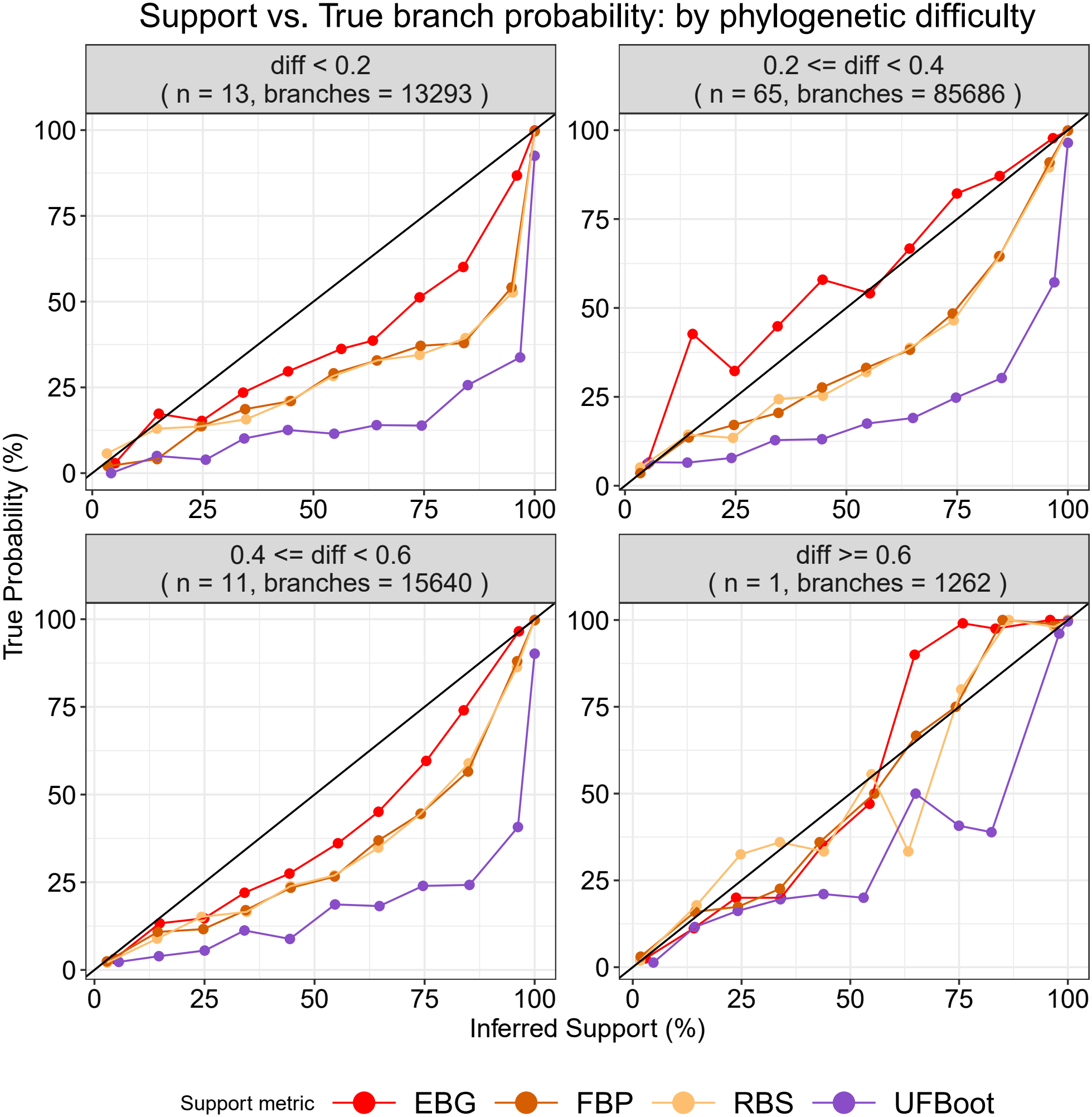
Support metric accuracy binned by phylogenetic difficulty.

**Fig. S10.**
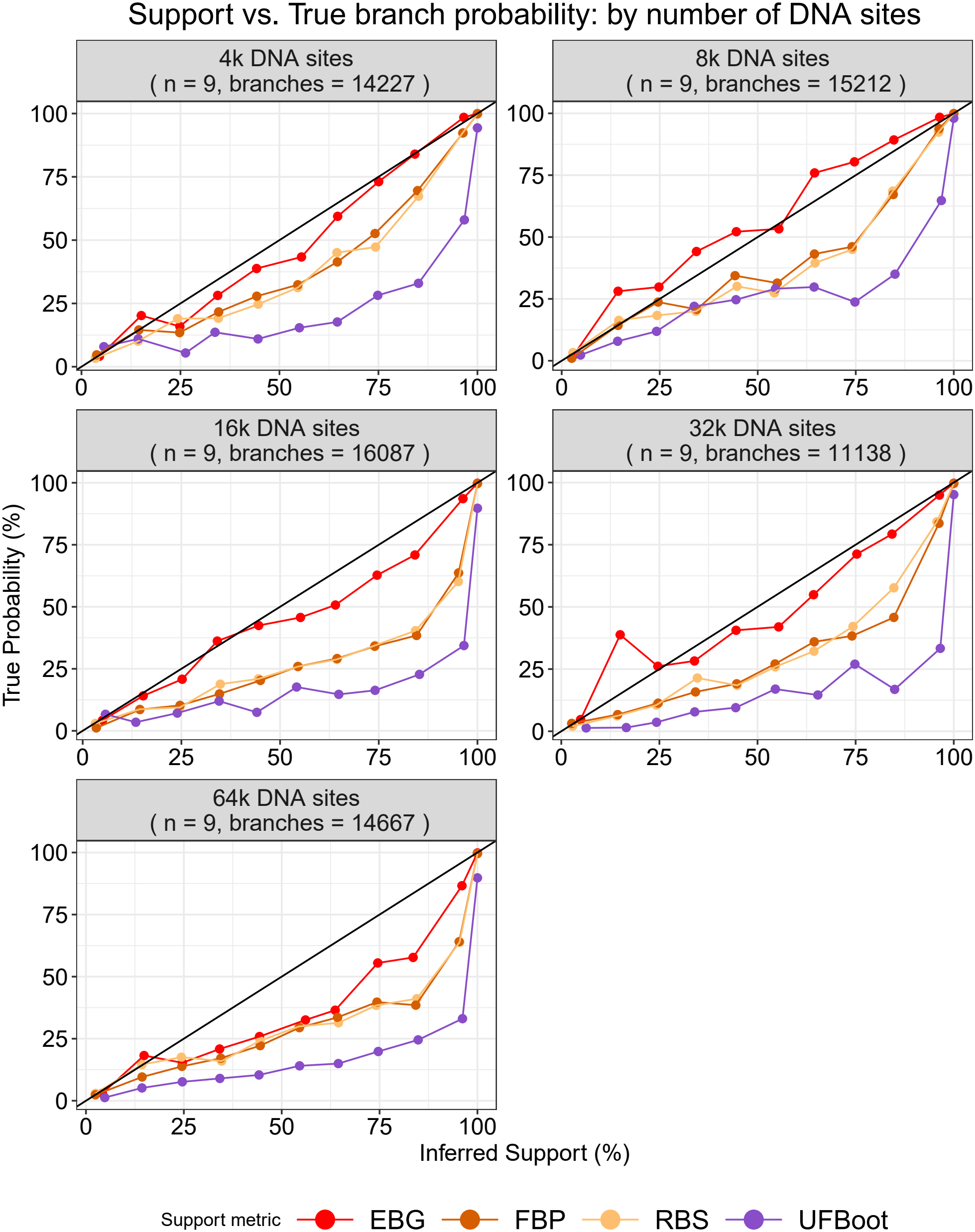
Support metric accuracy binned by number of DNA sites.

**Fig. S11.**
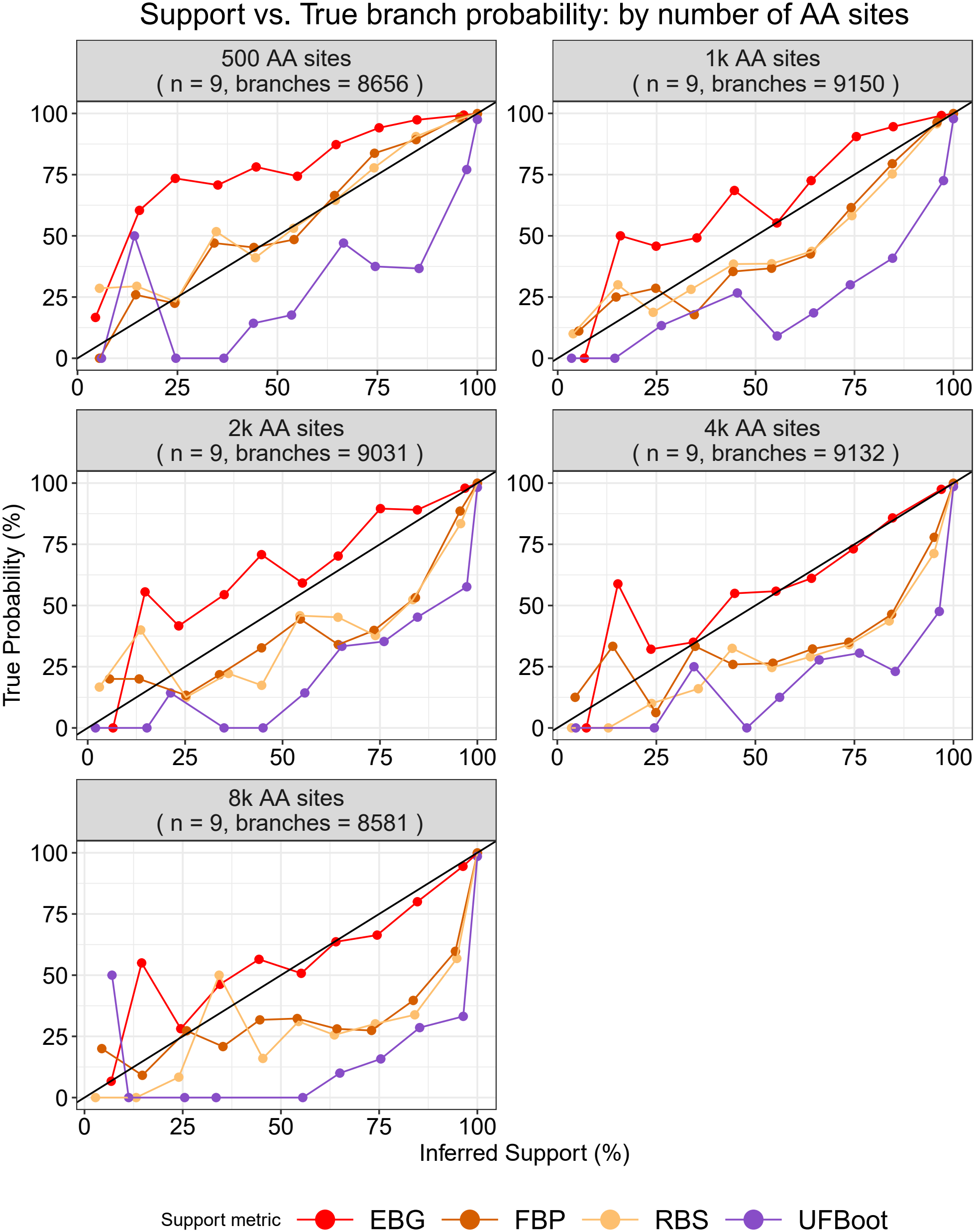
Support metric accuracy binned by number of AA sites.

